# Characterising the detectable and invisible fractions of genomic loci under balancing selection

**DOI:** 10.64898/2026.01.13.698512

**Authors:** Débora Y. C. Brandt, Aida M. Andrés, Tim Connallon, Max Reuter

**Affiliations:** Research Department of Genetics, Evolution and Environment, University College London, Gower Street, London WC1E 6BT, United Kingdom; UCL Genetics Institute, University College London, Gower Street, London WC1E 6BT, United Kingdom; Centre of Life Origins and Evolution, University College London, Gower Street, London WC1E 6BT, United Kingdom; School of Biological Sciences, Monash University, 25 Rainforest Walk, Clayton, Victoria 3800, Australia

## Abstract

Balancing selection refers to scenarios where selection maintains genetic polymorphisms affecting fitness and its components. Empirical studies have documented specific cases of balancing selection, yet the general prevalence of balanced polymorphisms within genomes remains a topic of longstanding debate. Although genome-wide scans for signals of balancing selection suggest that it is rare, current methods are notoriously conservative and may identify only a small and unrepresentative fraction of loci evolving under balancing selection. What, then, are the proportions of loci under balancing selection that are detectable versus invisible to genome scans? Here, we address this question using a combination of analytical modelling and population genetic simulations. Our results provide quantitative support for the intuition that a large fraction of loci under balancing selection will be invisible to standard tests for balancing selection, with the detectable loci representing a biased subset with large and symmetrical fitness effects on different fitness components. Quantifying power across parameters that vary between species also shows that the detectable and invisible fractions of genomic loci under balancing selection are likely to differ substantially between organisms. For example, a *Drosophila melanogaster*-like ratio of mutation to recombination rates (∼0.1) reduces detection power by roughly a quarter of the power expected with a human-like ratio (∼1). This, combined with other evidence suggesting that balancing selection could be common in *D. melanogaster*, showcases how our appreciation of the prevalence of balancing selection is limited by our ability to detect its genomic signals, especially in those species where it might be common.

## Introduction

Balancing selection occurs when natural selection actively favours the maintenance of genetic polymorphism. This can arise from several specific contexts of selection, including intrinsic fitness advantages of being heterozygous (*i*.*e*., ‘heterozygote advantage’ or fitness ‘overdominance’; (Fisher, 1922)), genetic trade-offs between different fitness contexts or environments (*e*.*g*., ‘antagonistic pleiotropy’ between fitness components: (Curtsinger et al., 1994); ‘sexually antagonistic’ trade-offs between sexes: (Kidwell et al., 1977)); trade-offs between temporal or spatially varying environments (Haldane & Jayakar, 1963; Levene, 1953), or negative-frequency-dependent selection, where genotypes become more favourable when rare (Fitzpatrick et al., 2007; Wright, 1942).

Loci under balancing selection are expected to be more polymorphic than loci evolving neutrally or under directional selection (though exceptions can occur when selection is weak or allele frequency equilibria under balancing selection are close to zero or one, (Robertson, 1962; Williamson et al., 2004)). When equilibrium frequencies are intermediate (*e*.*g*., very roughly between 0.2 and 0.8; (Hedrick, 1999)) and balancing selection is strong relative to genetic drift, the balanced polymorphisms should persist far longer than neutrally evolving polymorphisms (Nei & Roychoudhury, 1973), potentially over timescales spanning speciation events (Klein et al., 1998). Whatever their age, balanced polymorphisms should contribute disproportionately to standing genetic variation for fitness or fitness components (*e*.*g*., for survival, fertility, or longevity), with even modest numbers of such loci contributing substantially to fitness variation within populations (Charlesworth, 2015; Connallon & Czuppon, 2026; Crow, 1952, 1987; Zajitschek & Connallon, 2018).

The importance of balancing selection has, nevertheless, been a subject of longstanding debate in evolutionary biology. As an early and vociferous proponent, Dobzhansky (1955) argued that selection should often preserve genetic variation due to the greater versatility of heterozygous, relative to homozygous individuals in perpetually changing environments. Yet, by the close of the 20^th^ Century, high levels of molecular genetic diversity were reported by studies based first on allozymes (Lewontin & Hubby, 1966; Lewontin, 1974) and subsequently DNA sequences (Kreitman, 1983). The pervasiveness of these polymorphisms made Dobzhansky’s ‘balance hypothesis’ an implausible general explanation for molecular genomic diversity. Instead, the emerging neutral theory (Kimura, 1968; King & Jukes, 1969; Kimura, 1983) provided a simpler and more powerful explanation for the ubiquity of molecular genetic diversity, effectively sidelining balancing selection as irrelevant for all but a few exceptional loci within the genome (see p. 199 of Lewontin (1974); Hedrick (2012)).

More recent developments in evolutionary genetics have, however, prompted a resurgence in enthusiasm for balancing selection. Firstly, the coupling of field studies with genomics has identified exceptionally clear cases of balancing selection that maintains conspicuous phenotypic polymorphisms. Beyond classic examples like sickle cell anaemia (Allison, 1954), modern case studies include a pleiotropic trade-off between viability and mating success that maintains horn size polymorphism in Soay sheep (mapped to the *RXFP2* gene; Johnston et al. (2013)), a sexually antagonistic polymorphism in cuticular hydrocarbons of *Drosophila serrata* (in the *DsFAR2-B* gene; Rusuwa et al. (2022)), alleles affecting age of maturation in Atlantic salmon (the *VGLL3* gene; Barson et al. (2015)), temporally varying selection that maintains genetic polymorphism for insecticide resistance in *Drosophila melanogaster* (Karageorgi et al., 2025), and many others (reviewed in Bitarello *et al*. (2023); Ruzicka *et al*. (2025)). Second, the advent of high-throughput sequencing techniques and sophisticated analytical methods for detecting population genetic signatures of balancing selection has provided new means of identifying candidate loci under long-term balancing selection (reviewed in Bitarello et al., 2023; Fijarczyk & Babik, 2015). Applied to the human genome, such methods have provided strong evidence for long-term balancing selection in both African and European populations (Bitarello et al., 2018). Third, detailed lab studies of natural genetic variation in life-history traits suggest that *Drosophila* populations harbour higher levels of genetic variation than can be explained by models of mutation-selection balance (Charlesworth, 2015; Charlesworth & Hughes, 2000; Sharp & Agrawal, 2018). Balancing selection is a prime candidate to account for this apparent excess of genetic variation in life-history traits. If the same is true in other species, balancing selection may maintain substantial amounts of genetic and trait variation in natural populations.

While recent genome scans have identified putative balanced polymorphisms, it remains unclear how accurately they capture the true extent of balancing selection. The typically small fraction of loci with statistically significant signals of balancing selection (*e*.*g*., <1% of the genome in humans (Bitarello et al., 2018)) might reflect either the rarity of balancing selection across the genome or methodological limitations to detecting it. Most notably, classical methods cannot detect balancing selection unless intermediate-frequency polymorphisms are maintained over exceptionally long timescales. Loci segregating for evolutionarily ‘young’ balanced polymorphisms cannot easily be identified through population genomic approaches alone (but see Isildak et al. (2021) for a potentially promising approach), and old polymorphisms can likewise be missed due to limited data quality or other attributes that vary among species (*e*.*g*., mutation and recombination rates; effective population sizes) and may influence the strength of balancing selection signals. The balanced polymorphisms that are detected are, therefore, likely to represent only a minor, potentially unrepresentative, subset of loci evolving under balancing selection. In fact, a recent genome-wide analysis suggests there are hundreds of non-synonymous polymorphisms shared among human populations due to recent balancing selection, even though the method can only detect an aggregate signal across many loci rather than identify individual targets of balancing selection (Soni et al., 2022). While it remains an open question what proportion of these loci ultimately results in long-term stable balanced polymorphisms that are individually identifiable, this analysis suggests that at least recent episodes of balancing selection are widespread in the human genome.

Here, we aim to put the debate over the prevalence of balancing selection on a sounder conceptual and quantitative footing by evaluating factors affecting opportunities for long-term balancing selection and its detectability. It is obviously impossible to make general predictions about the proportion of a genome that is a potential target of balancing selection. Yet it is possible to assess—under a given model of balancing selection—how the population genetic parameters of genetic drift and selection will affect: *i*) the long-term maintenance of polymorphisms, and *ii*) the proportion of loci that develop signatures of balancing selection that are clear enough to be detectable using contemporary population genomic methods.

We address these questions while considering three mechanisms of balancing selection: overdominance, antagonistic pleiotropy and sexually antagonistic selection, each of which have been empirically documented in natural populations (e.g. Allison, 1954; Johnston et al., 2013; Rusuwa et al., 2022). These mechanisms represent a subset of balancing selection mechanisms (e.g. Glémin, 2021; Prout, 2000; Ruzicka et al., 2025), yet their evolutionary dynamics are representative of a broader array of balancing selection scenarios, which allows our predictions to be generalized. We begin by briefly reviewing predictions from population genetics theory, highlighting constraints associated with maintaining long-term stable polymorphisms under each mechanism of balancing selection. We then present simulations under each mechanism that explore the genomic signals associated with long-term balanced polymorphisms and assess how these signals translate into statistical power to infer the presence of balancing selection at these loci. Our analyses reveal how the maintenance of balanced polymorphisms and their detectability depend on the properties of the genetic polymorphism itself, as well as the genetic and demographic features of the species. They also provide a quantitative appreciation for the extent to which loci subject to balancing selection are detectable and suggest the biases in the specific types of balanced polymorphisms that are most and least likely to be identified.

## Results

### Long-term maintenance of balanced polymorphisms: Theoretical considerations

We start by reviewing theoretical predictions regarding the conditions under which genetic polymorphisms are stably maintained due to overdominant selection (OD), antagonistic pleiotropy (AP) or sexually antagonistic selection (SA). Our models focus on bi-allelic, diploid loci, and we assume that individuals mate at random and selection at each locus remains constant over time. Violations of these assumptions tend to restrict conditions for balancing selection (reviewed in Ruzicka et al. (2025), but see Wittmann et al. (2023) for signals of long-term balancing selection under temporally fluctuating selection). Full details of the models are presented in Appendix 1.

Long-term maintenance of a polymorphism under these models of selection requires two conditions. First, the combination of allelic fitness effects must actually generate balancing selection. Second, the resulting selection must be stronger than genetic drift for the polymorphism to be stably maintained over time. We deal with each condition in turn.

As far as the general criteria for balancing selection are concerned, OD is the most permissive scenario. In outbred populations, heterozygotes having higher fitness than both homozygotes (OD) invariably leads to balancing selection (Fisher, 1922). In contrast, AP and SA lead to balancing selection under only a subset of parameter conditions (Fig. 1). AP and SA each involve trade-offs between the fitness effects of alleles in different contexts of selection (between fitness components and between sexes, respectively). Considering a locus with alleles *A*_1_ and *A*_2_, *s*_1_ represents the homozygous cost associated with the *A*_1_ allele, *s*_2_ is the homozygous cost associated with the *A*_2_ allele, and dominance (*h*) defines the degree to which these costs manifest in heterozygotes (Fig. 1A). For simplicity, we focus on cases in which the dominance coefficient ranges from *h* = 0.5, implying co-dominance of the alleles, to *h* = 0, implying complete recessivity of these costs (for generalizations beyond these conditions, see Kidwell et al. (1977); Curtsinger et al. (1994)). Note that dominance values within the range 0 < *h* < 0.5 correspond to “dominance reversals” (Connallon & Chenoweth, 2019; Grieshop et al., 2024; Reid, 2022), where each allele is (partially) recessive in contexts where it is harmful and (partially) dominant in contexts where it is favourable.

**Figure 1.**
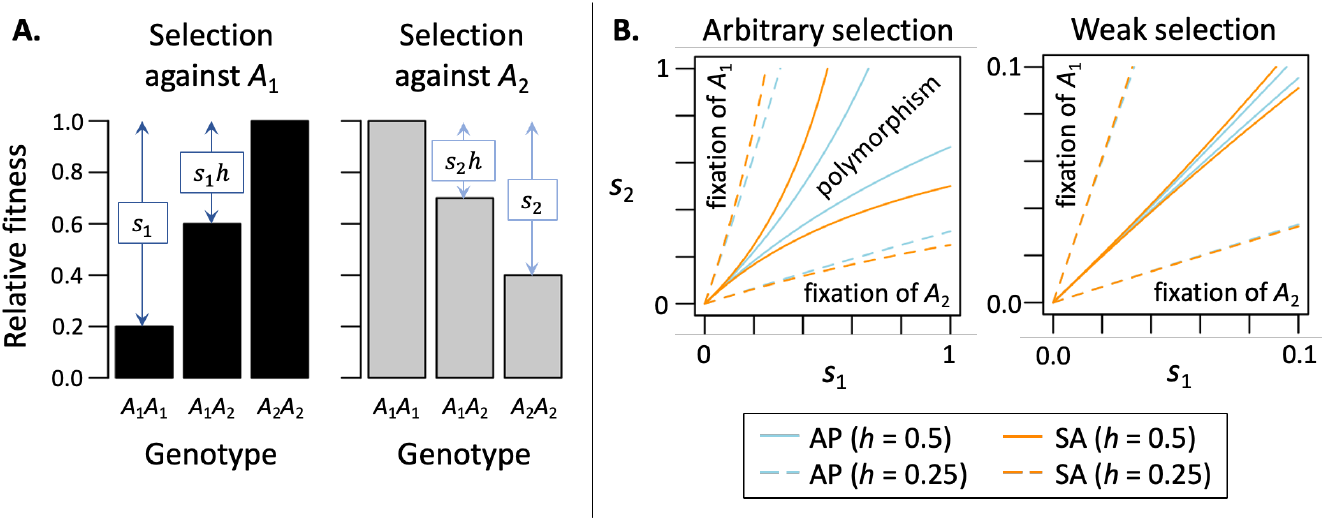
Conditions for balancing selection under antagonistic pleiotropy (AP) and sexually antagonistic selection (SA). **Panel A:** Visual summary of the AP and SA model parameters, with the example showing *s*_1_ = 0.8, *s*_2_ = 0.6, and *h* = 0.5. The two bar charts show the relative fitness of each of the three genotypes of a locus across two contexts of selection (i.e., different fitness components or sexes), where the *A*_1_ allele is selectively disfavoured in one context and the *A*_2_ allele is disfavoured in the other. **Panel B:** Criteria for balancing selection as a function of selection (*s*_1_ and *s*_2_) and dominance (*h*) parameters. The regions between each pair of curves (based on colour and line type) show conditions where selection favours the maintenance of polymorphism (i.e., where there is balancing selection). Predictions are shown when there is no dominance between alleles (i.e., alleles are co-dominant: *h* = 0.5), and when there is a beneficial reversal of dominance (*h* = 0.25), where negative allelic effects are partially recessive to beneficial effects. Panel B is modified from Connallon and Chenoweth (2019), with curves based on standard modelling results (see Kidwell et al. 1977; Curtsinger et al. 1994).

Balancing selection arises consistently in the SA and AP scenarios when the homozygous costs of each allele are symmetric in magnitude (*s*_1_ is roughly equal to *s*_2_), and the criteria for balancing selection expand with both the strength of selection and the strength of the dominance reversal (i.e., with *s*_1_ and *s*_2_ increasing and *h* decreasing; Fig. 1B). When selection is weak and there is no dominance reversal, only a small proportion of the parameter space will generate balancing selection. In this case, the proportion of random parameter combinations that leads to balancing selection is roughly 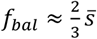 under AP and 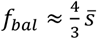 under SA, where 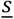 represents the average fitness effect (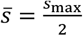 for parameters drawn from uniformly from the parameter space, with *s*_max_ representing the maximum selection coefficient; *s*_max_ ≪ 1 when selection is consistently weak). With dominance reversals (*h* < 0.5) and weak selection, the proportions expand to approximately 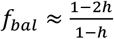 for both AP and SA, which corresponds to 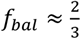 when *h* = 0.25 (as in Fig. 1B) and *f*_bal_ = 1 for a complete dominance reversal (*h* = 0).

The second condition for long-term stability of a balanced polymorphism is that selection towards a polymorphic equilibrium (which we denote as 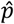) must be strong enough to prevent fixation of one or the other allele by genetic drift. The required strength of selection depends on the effective size of the population (*N*_e_) and the net strength of selection to the equilibrium (*α*), which differs between the mechanisms of balancing selection (*α* = *s*_1_ + *s*_2_ under the OD model, where *s*_1_ and *s*_2_ represent the fitness reductions of *A*_1_ and *A*_2_ homozygotes relative to the heterozygotes; *α* = (*s*_1_ + *s*_2_)(1 − 2*h*) + 2*s*_1_*s*_2_*h*^2^ and 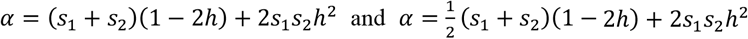 under the AP and SA models, respectively, with the SA and AP parameters, *s*_1+_, *s*_2_, and *h*, as depicted in Fig. 1A; see Appendix 2 for derivations of the equations for *α*).

The stationary distribution of the allele frequencies for a locus under balancing selection and drift (*i*.*e*., the predicted long-term probability distribution of allele frequency states) provides an intuitive measure of balancing selection’s ability to maintain polymorphism in the presence of genetic drift (Rice, 2004; Robertson, 1962). Balancing selection reliably maintains polymorphism when the allele frequency probability densities are much greater near the deterministic equilibrium (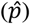) than near the boundaries (*i*.*e*., close to zero and one). Heuristically, balancing selection can be regarded as effectively strong in cases where the probability density around 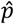 is at least ten times greater than the density near the boundaries. Under this definition, balancing selection is effectively strong when *α* > *a*_min_, where:

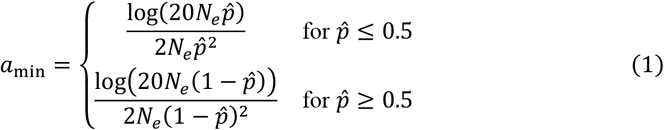

Figure 2A illustrates how the effective population size and equilibrium allele frequencies under balancing selection (*N*_*e*_ and 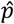) interact to determine criteria for effectively strong balancing selection (where *α* > *a*_min_). As can be seen in the figure, these conditions are most permissive in large populations with intermediate equilibria (note that *a*_min_ is always minimized at 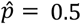).

**Figure 2.**
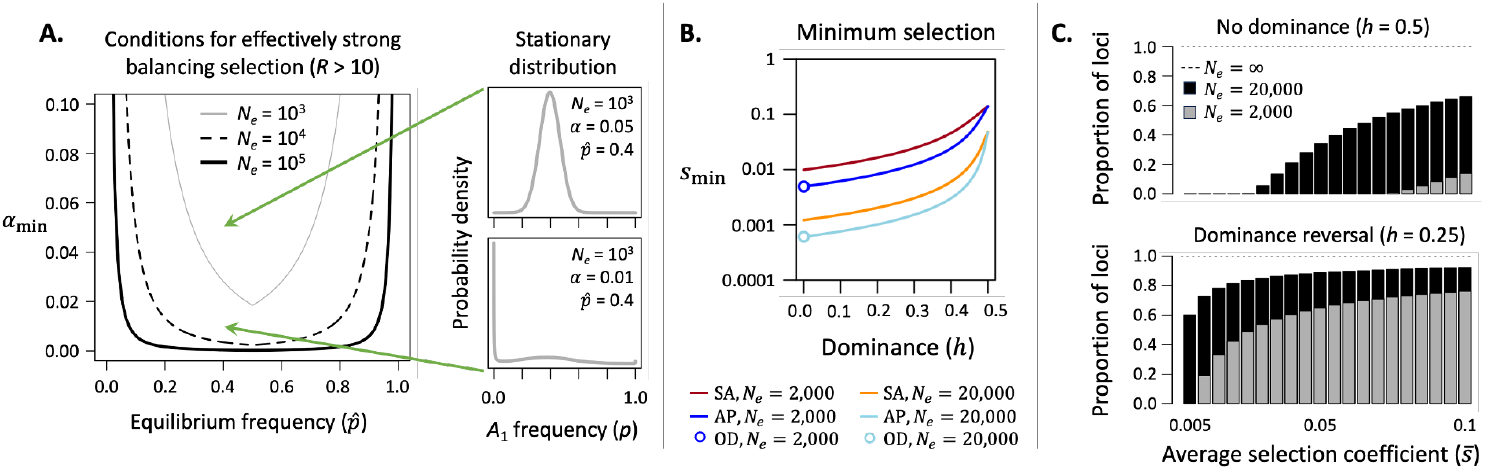
Effectively strong balancing selection. **Panel A** illustrates the thresholds for effectively strong balancing selection relative to genetic drift (*a*_min_; see eq. (1) and the surrounding text) for three population sizes (*N*_*e*_) and a spectrum of allele frequency equilibrium conditions under balancing selection 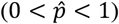. Regions above each curve denote combinations of *α* and 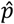 that generate effectively strong balancing selection (i.e., where the probability density for the stationary distribution is at least a 10-fold higher near the polymorphic equilibrium, 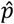, than near the allele frequency boundaries of zero and one, which correspond to fixation). Two special cases of the stationary distribution are shown: one where selection is ineffective (*α* < *a*_min_, and density is concentrated near *zero*) and one where selection is effective (*α* > *a*_min_, and density is concentrated near the equilibrium, 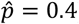). **Panel B** shows the minimum selection coefficients for each model that allow for effective balancing selection (the minimum is defined at 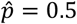, where conditions for effective balancing selection are most permissive). **Panel C** shows the proportion of loci under AP balancing selection that meet the criteria for effectively strong balancing selection. Results are based on randomly drawing 10^7^ sets of selection coefficients (*s*_1_ and *s*_2_) from independent uniform distributions between 0 and *s*_max_, where 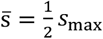 is the average selection coefficient. Bar heights were calculated as the number of parameter sets that met the criterion for effectively strong balancing selection 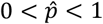, divided by the number of parameter sets that generated balancing selection 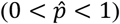.

Because *α* is defined differently for each model of selection (see above), thresholds for effectively strong balancing selection correspond to different absolute strengths of selection under the AP, SA, and OD scenarios. For example, the minimum values of *s*_1_ and *s*_2_ that are required for *α* > *a*_min_ (shown in Fig. 2B) are largest for the SA model, smallest for the OD model, and intermediate for the AP model (SA and AP cases are equal when *h* = 0.5, and AP and OD cases are equal when *h* = 0; see Fig. 2B).

A reduction of *N*_*e*_ amplifies drift and restricts the parameter space of effectively strong balancing selection (Fig. 2C), particularly so when dominance reversals are weak or absent. For example, when *h* = 0.5 and *N*_*e*_ = 2,000, SA and AP alleles cannot be reliably maintained polymorphic even by moderately strong selection (*e*.*g*., *s*_1_ ∼ *s*_2_ ∼ 0.05, corresponding to a 5% difference between the homozygote genotypes in each fitness component). Larger *N*_*e*_ and/or dominance reversals (*h* < 0.5) make long-term maintenance of polymorphism much more likely (Fig. 2C). Population genetic simulations (see Methods for simulation details) confirm these general predictions (Figs. S2, S4, S6). They further illustrate how the notion of maintaining long-term stable polymorphisms depends on the timescale of the observation: the range of parameter values that maintains polymorphism for 16*N*_*e*_ generations is significantly narrower than the range that maintains polymorphism for 8*N*_*e*_ generations (Fig. S2C,D).

### Empirical signatures of long-term balancing selection

The theory presented above summarizes factors that promote or constrain the maintenance of polymorphism at individual sites subject to balancing selection. Assessing the detectability of balancing selection requires additional considerations, because genomic scans for balancing selection do not focus on the balanced site itself but on signals of elevated genetic diversity at sites physically linked to the selected site. These signatures build up in the presence of long-term balancing selection, where the balanced polymorphism segregates for much longer than expected under neutrality (*i*.*e*., the time to the most recent common ancestor, TMRCA, at the selected site is greater than the neutral expectation of 4*N*_*e*_ generations). Genomic regions with an old TMRCA owing to balancing selection are expected to deviate from the genomic background in two ways. First, they are expected to show an elevated density of polymorphisms (and a concomitant deficit of substitutions relative to closely related outgroups) due to the long timespans over which mutations can accumulate since the TMRCA in the region and their lower probability of going to fixation due to linkage with the balanced site (DeGiorgio et al., 2014). Second, they are expected to show an excess of intermediate-frequency alleles when compared with the neutral expectation, as linked neutral variants tend to segregate at frequencies similar to those of the balanced polymorphism (see e.g., Fig. 1 in Bitarello et al. (2018)). These properties will be reflected in the local site-frequency spectrum (SFS) and provide the signal exploited by current population genomic methods for detecting long-term balancing selection (e.g. Bitarello et al., 2018; Cheng & DeGiorgio, 2020).

In the following section, we will explore how the presence and intensity of the signal, and thus our statistical power to identify regions under balancing selection, depends on the properties of the balanced polymorphism itself (*i*.*e*., the nature and strength of balancing selection, dominance, and the resulting equilibrium frequency of the balanced polymorphism), and factors that affect population genomic data, including the effective population size, and mutation and recombination rates, which vary among species and genomic regions.

We can get a preliminary sense of some of the key factors affecting neutral diversity near a long-term balanced polymorphism by quantifying the expected spike in heterozygosity near a balanced site under strong selection, which is analytically tractable. Consider a balanced polymorphism with alleles *A*_1_ and *A*_2_ maintained stably at deterministic equilibrium frequencies, 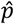 and 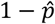, respectively. Building on previous work (see p. 103 in Rice (2004)), one can derive the expected width of the sequence surrounding the balanced site in which neutral polymorphisms will have an expected level of heterozygosity that is at least *K* times greater than the expected heterozygosity at unlinked neutral sites elsewhere in the genome (4*N*_*e*_*µ*, where *µ* is mutation rate per nucleotide site). The width of this region, in kilobases (kb), is:

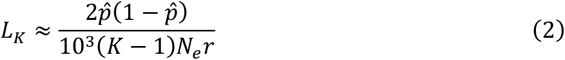

where *r* represents the recombination rate per nucleotide site (see Appendix 3). Inspection of eq. (2) shows that the width of elevated diversity surrounding the balanced polymorphism increases with the equilibrium heterozygosity of the selected site (with 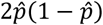), which is maximized when balanced alleles are maintained at equal frequencies 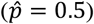, and it decreases with the population-scaled recombination rate per nucleotide site (*N*_*e*_*r*). Thus, we expect empirical signals of long-term balancing selection to be strongest when 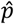 is intermediate and *N*_*e*_*r* is small, owing to small *N*_*e*_ and/or small *r*.

### Power to detect balancing selection in simulated data

While these predictions lend intuition, they ignore the reality that allele frequencies at the selected site fluctuate over time (*e*.*g*., due to drift), and overlook other empirically distinctive features of the SFS that are relevant to balancing selection scans, including the actual number of segregating linked variants and the shape of the SFS around a balanced site relative to the genome as a whole. Allele frequency fluctuations at the balanced polymorphic site are expected to reduce linked diversity relative to the more idealized case (as in eq. (2)) where the alleles are maintained at invariant equilibrium frequencies (for allele frequency fluctuations caused by temporally varying selection, see Wittmann et al. (2023)). Drift-mediated allele frequency fluctuations at the selected site should be minimal provided *N*_*e*_*α* is large and the equilibrium 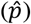 is intermediate, while fluctuations increase as *N*_*e*_*α* and 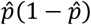 decline (Charlesworth & Charlesworth, 2010, pp. 354–355; Connallon & Czuppon, 2026).

To provide an assessment of statistical power for detecting balancing selection candidates, we turned to population genetic simulations. We ran extensive simulations (see Methods for details) with population-scaled values of *N*_*e*_*µ* and *N*_*e*_*r* chosen to roughly match those of humans and fruit flies—intensively studied species that may be representative of vertebrates and insects. We note that these simulations are not intended to provide specific power estimates for these species, since our models do not match their demographic histories (for example, the long-term *N*_*e*_ of human populations is smaller than 20,000 and their detailed demographic history generally increases power compared with the stable effective population size simulated here see Discussion). Rather, they are presented as toy examples that allow us to evaluate effects of relevant genomic parameters that vary among species. We first assessed power for a human-like ‘reference’ set of parameters (*N*_*e*_ = 20,000, *µ* = *r* = 10^-8^, age of polymorphism *T* = 8*N*_*e*_ =160,000 generations), which we will later use as a point of comparison with simulations using different population parameters. We assessed the power of the NCD method (Bitarello et al., 2018) to identify balancing selection in population genetic simulations of 10Kb chromosomes with a centrally located site under balancing selection and neutral variation in the flanking regions (see Methods). The results obtained for NCD should be representative of those obtained with other contemporary methods for detecting long-term balancing selection, which use similar approaches; additional results using the BalLeRMix method (Cheng & DeGiorgio, 2020) are presented in the Supplementary Figures S11 and S12 (and see Fig. S13 for a comparison between the two methods). NCD tests for an enrichment of polymorphism in the SFS by calculating, across all SNPs in a window, the average deviation of allele frequency from an intermediate ‘target frequency’. We used the equilibrium frequency of the balanced site as the target frequency for NCD, and our analyses therefore represent an idealised best-case scenario for power, where the true equilibrium frequency is known (Bitarello et al., 2018). For each set of parameters, we generated 100 replicate simulations in which the balanced polymorphism segregated until the end of the simulation (8*N*_*e*_ or 16*N*_*e*_ generations, thus conditioning on the maintenance of polymorphism; for results on the probability of maintenance over these time durations, see Figs. S1–S6). From these, we estimated power as the proportion of simulations that resulted in a statistically significant signal of balancing selection at P < 0.01, measured by NCD.

We evaluated the effects of our simulation parameters on power through multiple logistic regressions (Generalised Linear Models (GLM) with binomial error) with counts of successful and unsuccessful detection among the 100 trials of our simulations as dependent variables. The GLMs model power on the log-odds scale and allow us to estimate and test the effects of the simulation parameters while taking into account the fact that power is bounded between 0 and 1 as well as the binomial error of successful/unsuccessful detection of balancing selection (see Methods for more details).

In our simulations, the power to detect balancing selection is relatively low overall for our set of parameters, regardless of the exact method used for detection (i.e., NCD in Fig. 3 and Figs. S7–S9 versus BalLeRMix in Figs. S10–13). However, power varied considerably across parameter conditions in ways that broadly align with theoretical expectations. Among our human-like reference simulations, the variable that most strongly predicts power is the equilibrium frequency of the balanced polymorphism 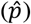, with the likelihood of detecting balancing selection increasing as the equilibrium approaches an intermediate frequency (i.e., 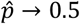; Fig. 3A). Across these reference simulations, the average power is 0.23 at 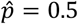, while power drops to 0.02 with 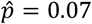. Fitting a binomial GLM with equilibrium frequency 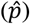, selection strength (*α*), selection model and their interactions as predictors of detection power, we found that the equilibrium frequency explains more than 60% of the variance in power across our simulations (Tab. 1). More specifically, the odds of successfully detecting balancing selection are estimated to increase by more than 30% with every 10% increase in the equilibrium frequency of the minor allele at the selected site, up to 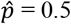 (b_eq. freq._ = 2.737 on log-odds scale, estimated for OD).

**Table 1.** Binomial GLM of numbers of instances of successful/unsuccessful detections of balancing selection in the reference set of simulations, as a function of equilibrium frequency 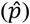, selection strength (*α*), selection model and their interactions. Only terms with P-value <0.05 are shown.

| Predictor | Deviance | % deviance explained | P-value |
| --- | --- | --- | --- |
| $\hat{p}$ | 362.96 | 62.28% | < 0.001 |
| $\alpha$ | 47.66 | 8.2% | < 0.001 |
| $\alpha * \hat{p}$ | 13.64 | 2.3% | < 0.001 |
| $\hat{p} * \text{selection model}$ | 13.68 | 2.4% | 0.001 |
| $\alpha * \hat{p} * \text{selection model}$ | 33.38 | 5.7% | < 0.001 |

**Figure 3.**
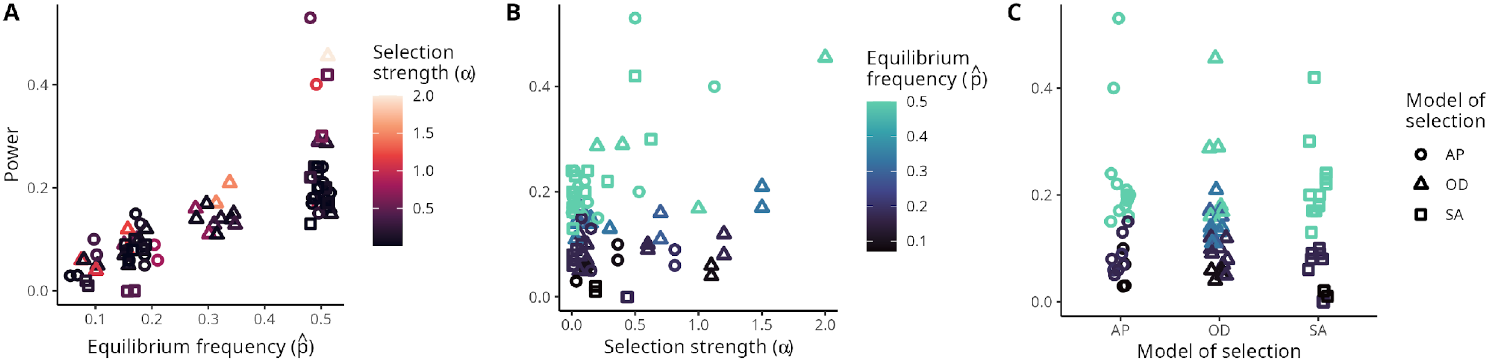
Power as a function of equilibrium frequency (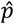, panel A), selection strength (*α*, panel B), and model of selection (panel C) for the reference simulations. On panels A and C points were horizontally jittered for visualization purposes. The three panels show the same data points, with power on the vertical axis plotted against different horizontal axes of variation.

Power also generally increases with the strength of selection (Tab. 1), but this effect depends on interactions with other parameters. First, the positive effect of selection strength on power is pronounced at intermediate equilibrium frequencies but declines and eventually vanishes when equilibrium frequencies of the minor allele at the selected site are lower (Tab. 1, Fig. 3B). Second, this interaction effect differs between the three models of selection and is strongest under SA. Under OD and AP, increasing selection strength by 0.1 increases the odds of detecting balancing selection by 5.7% and 1.6% for equilibrium frequencies of 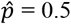 and 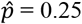, respectively. For SA, in contrast, the same change in selection strength increases the odds of detection by 15.7% for an equilibrium frequency of 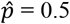 but decreases the odds by 40% for an equilibrium of 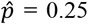 (see Fig. 3B for simulation data and Fig. 4 for model predictions). Beyond these interactive effects, however, the model of selection does not impact the overall power of detecting balancing selection (Tab. 1, Fig. 3C).

**Figure 4.**
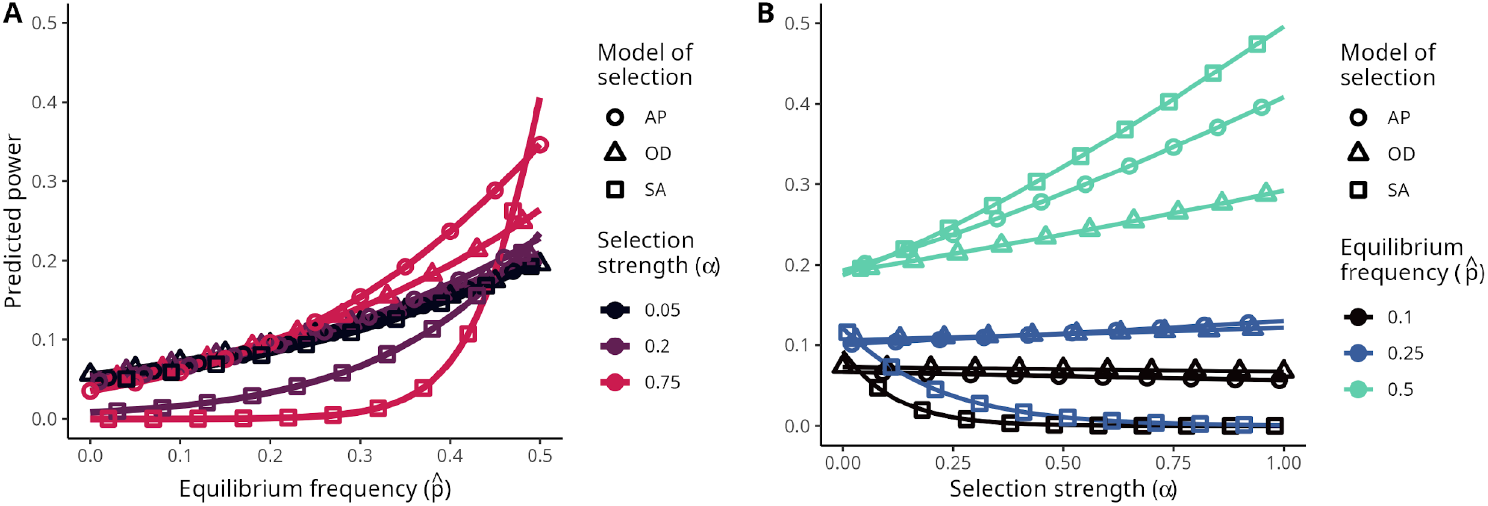
Power predicted by the Generalised Linear Model fitted to the data for standard simulations as a function of equilibrium frequency (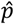, panel A) and selection strength (*α*, panel B). Results are shown for different parameter combinations as indicated by the right-hand legend of each panel.

As current methods for inferring targets of long-term balancing selection rely largely on a signal of excess polymorphism and intermediate frequencies in the SFS, power should increase in conditions that generate a larger and wider signature of excess polymorphism and decrease under those that restrict the signal. To explore these effects, we ran additional simulations under OD while varying the recombination and mutation rates, the effective population size, and the age of the balanced polymorphism. These simulations show that power consistently decreases at higher recombination rates (Tab. 2), particularly so in cases where the equilibrium frequency is intermediate (Fig. 5A; Tab. 2). The model predicts that a ten-fold increase in recombination rate decreases the odds of detection by roughly ten-fold when the equilibrium is intermediate 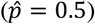, but only by 4.5-fold for an equilibrium of 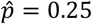 (Tab. 2, b values).

**Table 2.** Binomial GLM of numbers of instances of successful/unsuccessful detections of balancing selection (overdominance model) as a function of predictors and interactions. Only terms including the variable of interest and with P-value <0.05 are shown.

| Predictors included in the model | Predictors discussed in main text | Deviance | % deviance explained | P-value | b (odds ratio) |
| --- | --- | --- | --- | --- | --- |
| equilibrium frequency ($\hat{p}$), selection strength ($\alpha$), $\log_{10}$ of **recombination rate (r)** | r | 249.89 | 55.3% | <0.001 | -0.841 (0.43) |
| | $r * \hat{p}$ | 11.12 | 2.5% | 0.001 | -2.766 (0.06) |
| equilibrium frequency ($\hat{p}$), selection strength ($\alpha$), $\log_{10}$ of **mutation rate (m)** | m | 112.86 | 18.9% | <0.001 | 0.879 (2.41) |
| | $m * \hat{p} * \alpha$ | 3.98 | 0.7% | 0.046 | 1.776 |
| equilibrium frequency ($\hat{p}$), selection strength ($\alpha$), **allelic age (t)** | t | 557.1 | 51.2% | <0.001 | 0.002 (1.002) |
| | $t * \hat{p}$ | 69.17 | 6.4% | <0.001 | 0.015 (1.015) |
| | $t * \alpha$ | 12.50 | 1.1% | <0.001 | -0.002 |
| equilibrium frequency ($\hat{p}$), selection strength ($\alpha$), **effective population size ($N_e$)** in thousands of generations | $N_e$ | 82.02 | 23.2% | <0.001 | -0.036 (0.96) |
| | $N_e * \hat{p}$ | 17.92 | 5.1% | <0.001 | 0.033 (1.03) |
| | $N_e * \hat{p} * \alpha$ | 6.51 | 1.8% | 0.011 | 0.130 (1.14) |

**Figure 5.**
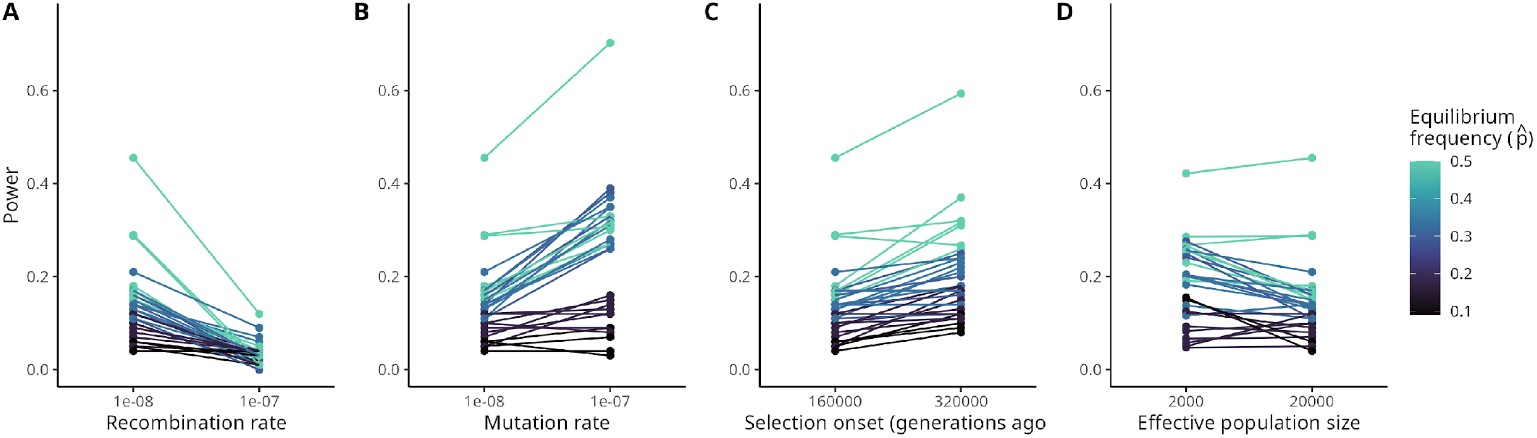
Power to detect balancing selection in our reference set of simulations (points on the left of panels A-C and on the right of panel D) compared to simulations with increased recombination rate, mutation rate, age of the selected allele and reduced effective population size. Balancing selection was simulated under overdominance, and parameters in the reference set of simulations are *N*_*e*_ = 20,000, recombination and mutation rates of 10^-8^ per site per generation, selection onset time 160,000 generations ago.

Conversely, power increases with a higher mutation rate per nucleotide site (Fig. 5B). The model predicts that the odds of successfully detecting balancing selection increase by about 150% for each ten-fold increase in mutation rate (Tab. 2, b value). Note that while there is no interaction of mutation rate with the allele frequency equilibrium (Tab. 2), there is a nominally significant triple interaction with the equilibrium frequency and strength of selection, though this explains a negligible proportion of variation in power (Tab. 2).

The intensity of the signal of balancing selection also depends on the timespan over which a balanced polymorphism has been maintained, with the age of the balanced polymorphism in our simulation data having a strong positive effect on our power to detect it (Fig. 5C; Tab. 2). In our analyses, the allelic age also interacts significantly with the equilibrium frequency in determining power (Tab. 2). Accordingly, for every additional 0.5 *N*_*e*_ generations of allelic age in our model, the odds of successfully detecting balancing selection increase by 9.5% for a polymorphism with an equilibrium frequency of 0.5 and by 5.5% with an equilibrium frequency of 0.25.

Finally, increasing the effective population size generally reduces the detectability of balanced polymorphisms, though effects of *N*_*e*_ are complex (Fig. 5D). The binomial GLM reveals a significant negative effect of *N*_*e*_ on power (Tab. 2), with a significant interaction with the equilibrium frequency and with the interaction of equilibrium frequency with selection strength (Tab. 2). Thus, the GLM predicts that among loci with a minor allele equilibrium frequency close to our lower bound of 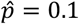, and under weak selection (*α* = 0.1), a ten-fold increase in *N*_*e*_ lowers the odds of successfully detecting balancing selection by over 30%. In contrast, for loci with an intermediate equilibrium frequency of 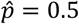 and under strong selection (*α* = 1.5), the GLM predicts that the same increase in *N*_*e*_ would reduce the odds of detection by only 10%.

## Discussion

We assessed the propensity of balancing selection to maintain long-term stable polymorphism and the power of current inference methods to successfully detect such loci. The main message from our analyses is that detecting balanced sites in population genomic data can be difficult, at least in the populations of constant size that we simulate here. Even where balancing selection is strong relative to drift and has maintained allelic variation at frequencies closely clustered around an equilibrium frequency of 0.5 for time spans that vastly exceed neutral coalescent times (such as 16*N*_*e*_ generations), state-of-the-art methods of inference still only rarely exceed a power of 50%. For example, a mutation under OD with relatively strong and symmetrical selection (*s*_1_ = *s*_2_ = 0.1, resulting in 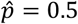) that is 8*N*_*e*_ generations old is detected as a balanced locus with a probability of only 28% in our reference simulations. These selection coefficients are of the same order of magnitude as the ones estimated for the remarkable example of net overdominance maintaining horn size polymorphism in Soay sheep (Johnston et al. (2013), see their Tab. S3). Our results thus show that in our models, 72% of loci under similar selection strengths and equilibrium frequencies would be missed. This finding contrasts with the notion that inference power with current approaches is good (Bitarello et al., 2023), a perception that is probably due to the empirical focus on human populations, which have particular demographic histories that are generally expected to boost the power of existing methods (see below).

While power in our simulations is moderate in the best of cases, it further decreases, often substantially, under less ideal and perhaps more realistic conditions. The selection parameters have particularly pronounced effects on the long-term maintenance and eventual detectability of a balanced locus. Selection and dominance coefficients determine whether a polymorphism will be subject to balancing selection at all (as in the AP and SA cases), the equilibrium frequencies of balanced polymorphisms, and the efficacy with which selection maintains the alleles near equilibrium. Each of these factors influences the long-term stability of a balanced polymorphism. But even when a polymorphism is long-term stable, balancing selection resulting in allele frequency equilibria close to zero or one are exceedingly unlikely to be identified, not only because they are more susceptible to becoming lost by drift, but also because even those polymorphisms that persist produce weak signals of balancing selection that are very hard to identify.

### Model of selection mainly affects maintenance

The models of selection differ very strongly in the strength of selection they generate and hence their propensity to maintain polymorphism in the first place. With OD the net strength of selection is a simple linear function of its underlying selection coefficients (*α* = *s*_1_ + *s*_2_), leading to effectively strong net selection in cases where population-scaled selection parameters are large (i.e., *N*_*e*_*s*_1_ ≫ 1 and *N*_*e*_*s*_2_ ≫ 1). In contrast, for scenarios of AP and SA without dominance reversals, the net strength of balancing selection (when it occurs) depends on the product of the selection coefficients (i.e., 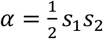 when *h* = 0.5), so that the same values of *s*_1_ and *s*_2_ lead to much weaker balancing selection under AP and SA than under OD (see Appendix 2 for derivations of equations for *α*). Assuming the distribution of fitness effects is similar under the three mechanisms, then those subject to antagonistic selection should be less likely to remain polymorphic over extended periods of time and, where they do, be harder to detect by genome scans for balancing selection. While dominance reversals dampen these differences between the antagonistic selection scenarios and OD, balancing selection in the SA and AP scenarios are expected to remain weaker and less permissive than the OD case unless dominance reversals are complete.

While the specific model of balancing selection has a strong effect on the propensity to maintain balanced polymorphisms, it did not have an overall effect on detection power. However, our analyses revealed more subtle interactions between the model of selection, the net strength of selection (*α*), and the equilibrium frequency 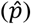. Specifically, at low equilibrium frequencies, power decreases with increasing strength of selection under SA, while selection strength has little or no effect on power under OD and AP. This effect is surprising, given that *α* is intended to capture the strength of selection in a way that is independent of the mechanism generating it. We note, however, that the expressions for *α* are approximations based on the assumption of effectively weak selection (see Appendix 1). It is therefore possible that the interaction effects we detected here were due to higher-order terms that are not included in the calculation of *α*.

### Selection onset and effective population size

Our simulations also demonstrate that the probability of inferring balancing selection at a locus increases with the age of the polymorphism, as signatures of linked variation build up over time. Importantly, the timescale of this process increases with the effective size of the population. For the signature of selection in linked variation to be discernible, the coalescent time at the balanced site needs to substantially exceed the average neutral coalescent time of 4*N*_*e*_ generations. This means that when *N*_*e*_ is larger, balanced polymorphisms need to be older, in generations, to be detectable. Likewise, for a given age in generations, polymorphisms are harder to detect in species with high *N*_*e*_ than in those with low *N*_*e*_.

But the timescales over which signatures of selection accumulate, and how they differ between species, also matter for the presence of balancing selection more generally. Thus, the time for signatures to build up can be considerable in geological time. In humans, for example, an age of 8*N*_*e*_ generations corresponds to roughly 2My (based on a historical *N*_*e*_ of 10^4^ and a 25-year generation time), or roughly a quarter of the time since the human–chimpanzee split. For a polymorphism to be present and detectable after this time, sizable fitness effects need to persist for this duration. This could substantially limit the prospects for the presence of long-term balancing selection in species with large *N*_*e*_, since stability of the polymorphism requires a relative constancy of the environment over a long time. However, this effect is attenuated by the fact that generation times tend to correlate negatively with *N*_*e*_. An analysis of the relationship between *N*_*e*_ and generation time (*g*) (Chao & Carr, 1993) suggests that, very roughly, 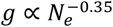, in which case the geological timescale (in years) that is required to generate a genomic signal of long-term balancing selection is on the order of 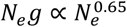. Thus, despite the inverse relationship between *N*_*e*_ and *g*, the timescale for generating a signal of balancing selection should generally be higher in lineages with larger *N*_*e*_ and shorter *g* (such as insects). As a consequence, for a given geological age of a polymorphism, the signals of balancing selection are less likely to be visible in large-*N*_*e*_ lineages because the criteria for “long-term balancing selection” (as defined in Bitarello et al., 2023) are more restrictive.

Interestingly, *Drosophila melanogaster* may be an exception to the broader pattern, in that it has a short generation time (∼15 generations/year or 0.07 years/generation, (Turelli & Hoffmann, 1991)) but a relatively modest effective population size (i.e., historical *N*_*e*_ of roughly 10^6^ (Arguello et al., 2019)). Based on these figures, 8*N*_*e*_ would correspond to 0.56My, much shorter than the equivalent time in humans, which makes it more likely that broader environmental conditions remain constant enough to allow for the maintenance of balanced polymorphisms in this species. In addition, extremely short generation times such as those in *Drosophila* provide further scope for balancing selection due to seasonal fluctuations in the environment (Wittmann et al., 2023), which is an unlikely avenue for balancing selection in species with long generation times.

The effective population size also has other direct and indirect effects on the presence and detectability of balanced polymorphisms, beyond the timescales discussed above. Large *N*_*e*_ leads to reduced drift and concomitantly more effective selection to maintain balanced polymorphisms and should therefore (all else being equal), increase the proportion of balanced loci that remains long-term polymorphic. On the other hand, *N*_*e*_ also has adverse effects on the signatures at linked neutral polymorphism. Species with large *N*_*e*_ tend to have larger effective recombination rates (larger *N*_*e*_*r*), owing trivially to the effect of larger *N*_*e*_ but also due to the fact that large-*N*_*e*_ species often have relatively streamlined genomes and accordingly higher per-nucleotide recombination rates (*r*) (Lynch, 2006). Elevated recombination will erode the balancing selection signal of intermediate frequencies at neutral sites that are closely linked to selected variants and thereby make the signature sparser and inference harder. This effect can be compensated by a higher mutation rate (*µ*), which favours the generation of linked polymorphism around the balanced site. Thus, for a given *N*_*e*_, the power to detect balancing selection is determined by the ratio of mutation to recombination rate, *µ*/*r*.

### Demographic history and comparison to previous power estimates

It is worth noting that while we have only simulated simple scenarios of constant population size, more complex demographic histories are likely to substantially affect detection of balanced polymorphisms. Changes in the effective population size over time, such as sudden contractions (bottlenecks) and subsequent expansions, can have beneficial effects on detection. Power can be improved because a bottleneck in the not-too-distant past will result in increased rates of coalescence (a more recent average TMRCA) at neutral regions that are unlinked to sites under balancing selection, and thus on average tend to cause a leftward-shifted SFS at those regions. Peaks of intermediate frequencies and older TMRCAs at sites linked to balanced loci stand out more clearly from such genomic backgrounds of neutral diversity. It is likely that this phenomenon has facilitated the identification of balanced loci in humans, where migration out of Africa has resulted in a series of bottlenecks and founder events that have systematically shifted the neutral SFS.

The small *N*_*e*_ and specific demographic history of humans likely explains why previous analyses of the power of current inference methods, including the NCD statistic used here (Bitarello et al., 2018) or BalLeRMix (Cheng & DeGiorgio, 2020), have provided higher estimates of power (reaching over 80% under some conditions) than the ones we report here for our human-like ‘reference’ parameters. While methods like NCD or BalLeRMix are general in scope, they have typically been benchmarked using simulated data that follow a demographic history and a mutation-to-recombination rate ratio that is characteristic of human populations (e.g., 2.5 in Bitarello et al. (2018) and sampled between 2.5 and 0.83 in Cheng and DeGiorgio (2020)), and relatively old polymorphisms (*e*.*g*. 5 million years in Bitarello et al. (2018)and up to 650,000 generations with *N*_*e*_ = 10^4^ in Cheng and DeGiorgio (2020), which corresponds to 16.25 million years with the generation time of 25 years used in Bitarello et al. (2018)). Given the specific assumptions made in the original method publications and the substantial variation in power as a function of the parameters used, we reiterate that it is imperative to combine any application of existing methods to a careful power analysis, parameterised as much as possible to fit the study organism and population (Bitarello et al., 2018, 2023).

### Methodological notes on NCD

We have deliberately focused our analyses on a single approach, NCD (but see Supplementary Material for additional analyses with BalLeRMix), to evaluate how different population genetic parameters determine the efficacy of balancing selection and the signals it generates. It is not our aim to compare different methods, all of which essentially exploit the same signals in the data and will hence be affected similarly by the parameters we explore. Nevertheless, we will briefly discuss a few specificities of our application of NCD that are expected to have small effects on the absolute power we estimate.

First, in our analyses, we use the actual equilibrium frequency, which is known in our simulations, as the NCD ‘target frequency’ (i.e., the frequency value from which NCD measures the mean squared distance of allele frequencies in a window). Doing so will have a small positive effect on power, compared to the case of an empirical application where the target frequency is not known, and is therefore set to an arbitrary intermediate value. Previous analyses (Bitarello et al., 2018), however, have shown that power is minimally impacted by the exact frequency value of the target and additional analyses performed on our simulations confirm this (Fig. S14).

On the other hand, two other specificities of our application of NCD are likely to have a small negative effect on power. First, we use a variant of NCD, NCD2, that takes into account the SFS of derived alleles, including substitutions. Because we do not simulate an outgroup, however, NCD2 is not as powerful here as the approach would be in a real-world application where the SFS also includes fixed sites that result from substitutions inferred from outgroup data (see Methods). Second, we use NCD with a fixed window size of 3Kb, independently of the recombination rate. In cases where recombination rate is high and regions of elevated polymorphism are narrow, calculating the SFS over a smaller interval can improve power, although this is offset by the fact that the small numbers of SNPs in a narrow focal window generate noise that negatively impacts the inference (Fig. S15). Again, real-world applications should be preceded by power analyses that explore these effects and can help to optimise the analysis.

### General considerations on differences between species

Our analyses make it clear that the prospects of identifying targets of balancing selection are expected to vary significantly and systematically among species. Thus, species will harbour variable numbers of long-term balanced loci, depending on the effective sizes of their populations. Furthermore, the effective size, combined with mutation and recombination rates will determine how difficult these sites are to identify as targets of balancing selection. In humans, for example, the small effective population size means that only a limited number of balanced polymorphisms are expected to have remained polymorphic over sufficiently long periods of time. On the other hand, the same low effective population size, together with the high ratio of mutation to recombination rate (*μ*/*r* = 1.2 × 10^−6^/1.3 × 10^−6^∼1 (Pritchard, 2023 Chap. 1.1)), favour detection of such loci. In contrast, *D. melanogaster*, and possibly other insects, presents a much harder case for identification, as the recombination rate is high relative to the mutation rate (*μ*/*r* ∼ 3 × 10^−7^/3 × 10^−6^ = 0.1 (Wang et al., 2023)), which our analyses suggest should reduce power by roughly four-fold compared to a species with human-like mutation and recombination parameters (mean power across simulated parameters in Fig. 4A, 0.14 vs. 0.03).

Of course, as discussed above, there are other important factors at work that affect the relative frequency of detectable balanced loci in the two species, such as the time required for long-term balancing selection, the scope for seasonal selection, effects of *N*_*e*_ on the maintenance of polymorphism on the one hand and its detectability on the other, and species-specific demographic histories. Overall, there is little doubt that the detection of balancing selection in *Drosophila* is challenging, which suggests that there could be many targets of balancing selection in *D. melanogaster* that are invisible to genomic scans for balancing selection, and this may also be the case for many other species with similar population parameters. Indeed, a genome-wide screen for balancing selection in fruit flies combining Theta and Tajima’s *D* identified roughly 100 candidate balanced loci (Croze et al., 2017), compared to over 1000 candidates in humans (Bitarello et al., 2018). Although more sophisticated methods may improve power in *Drosophila*, the notion that large numbers of balanced polymorphisms might exist and be missed in genomic scans is in line with the high levels of standing genetic variation for major fitness components in this species, which exceeds the variation explainable under mutation-selection-drift balance (Charlesworth, 2015; Sharp & Agrawal, 2018). The excess variance in life-history traits might be attributable to balanced polymorphisms segregating in *D. melanogaster* populations. While it seems plausible that at least some of these balanced polymorphisms would be detected if detection methods were more powerful, others could remain invisible if they are relatively young (relative to 4*N*_*e*_). It is difficult to assess these alternatives in the absence of more powerful approaches, and similar conclusions may apply to many other species with equally unfavourable population parameters.

## Methods

### Population genetic simulations of balancing selection

We used simulations to assess which regions of the parameter space generate genomic signatures of balancing selection that can be detected using current approaches. Simulations were run in SLiM (version 4.0.1, (Haller et al., 2019)), a forward simulation framework for population genetics. In line with the analytical derivations above, we implemented overdominance (OD), antagonistic pleiotropy (AP) and sexually antagonistic selection (SA), in addition to simulations under neutrality. In the simulations with selection, we simulated a single focal site, embedded centrally in a chromosome of 10,000bp, large enough for neutral sites in the periphery to be unaffected by selection at the central site. To implement the three selection mechanisms, we used custom functions (see SLiM scripts in available in https://github.com/reuterlab/bls_sim/) to run a non-Wright-Fisher model for populations of constant size with non-overlapping generations. In each generation, the code generated offspring by randomly drawing parents based on their fitness values, calculated according to their genotype at the locus under selection and the selection mechanism and parameters (selection coefficients, dominance, cf. Tab. A1 in Appendix 1) being simulated. We used the same code for reproduction in the neutral simulations, but set the fitness of all individuals to 1. In all simulations the default sex ratio of 1:1 was used.

At the start of each simulation, a mutation at the focal site was introduced on a single chromosome, randomly chosen within the population. The mutation would be of type A_1_ or A_2_ (see Tab. A1, *e*.*g*., either male- or female-beneficial for SA) with equal probability. To reduce the number of simulation runs where the polymorphism was quickly lost due to drift when rare, we initially subjected the mutant allele to directional positive selection with *s* = 0.5 and *h* = 1/2 until the mutant allele frequency reached 5% (or the mutant allele was lost). This selective boost facilitates the establishment of the polymorphism, but due to its old age and short duration should have negligible influence on the patterns of linked variation. For polymorphisms that reached a frequency of 5%, simulations then switched from directional to balancing selection with a specified mechanism and parameter combination. We recorded the frequency of the mutant (derived) allele and the genealogy of all chromosomes in the population at 16000, 32000, 160000 and 320000 generations.

To optimise the runtime of simulations, we did not include neutral mutations in any of the SLiM simulations. Instead, we recorded the full chromosome-wide genealogies from each simulation (using ‘tree-sequence recording’, (Haller et al., 2019)). We then used ‘recapitation’ in pyslim (version 1.0.4 (Ralph et al., 2023)) to generate the preceding genealogical relationships of the starting chromosomes up to their most recent common ancestor. Finally, we added neutral mutations on the complete recapitated genealogies with msprime (version 1.2.0, (Baumdicker et al., 2022)).

The parameter values used in our simulations are summarised in Table 3. We explored selection and dominance coefficients to match the analytical results. For the species-specific parameters of effective population size (*N*_*e*_), recombination rate (r) and mutation rate (µ) were chosen to cover a broad range of relevant scenarios. Thus, we simulated population sizes of *N* = *N*_*e*_ = 2,000 and *N* = *N*_*e*_ = 20,000 and neutral mutation at rates of µ = 10^-7^ and µ = 10^-8^ per base per generation, such that the expected genetic diversity (*π* = 4*N*.*μ*) ranges from 8×10^-5^ to 8×10^-3^. The highest expected genetic diversity (for *N*_*e*_ = 20,000 and µ = 10^-7^) approaches that of *Drosophila melanogaster* (*π* = 10^-2^ (Leffler et al., 2012)), while the combination of *N*. = 20,000 and µ = 10^-8^ generates diversity similar to that estimated in humans (*π* = 8×10^-4^ (Pritchard, 2023 Chap.1.1)). We simulated recombination rates of r=10^-7^ and *r* = 10^-8^ per base per generation to include combinations of µ = *r* =10^-8^, which are similar to the values estimated in humans (Pritchard, 2023 Chap. 1.1), as well as a ratio of mutation to recombination rate similar to that estimated in *Drosophila melanogaster* (*µ*/*r* = 10^-8^/10^-7^ = 0.1, (Wang et al., 2023)) and a scenario where mutation rate exceeds recombination rate (*µ*/*r* = 10^-7^/10^-8^ = 10), which should facilitate the detection of signatures of selection. Finally, we used two values for the onset of selection (or age of the polymorphism), *T* = 8*N*_*e*_ and *T* = 16*N*_*e*_ (16,000 and 32,000 generations for *N*_*e*_ = 2,000 and 160,000 and 320,000 generations for *N*_*e*_ = 20,000). These values roughly correspond to the definitions of long-term and ultra long-term balancing selection (Bitarello et al. 2023). Both significantly exceed the expected coalescence time (time to the most recent common ancestor, TMRCA) under neutrality, 4*N*_*e*_. The probability of observing a TMRCA larger than 8*N*_*e*_ generations under neutrality is approximately 0.054, while that of a TMRCA larger than 16*N*_*e*_ generations is approximately 0.001 (Wakeley 2009, p.77 eqn 3.27).

**Table 3.** Parameter values used in the simulations. We highlight in bold the values used in our “reference” set of simulations, which approximate the parameter values estimated in the literature for humans.

|  |  |
| --- | --- |
| Selection model | Overdominance (OD)<br>Antagonistic pleiotropy (AP)<br>Sexual antagonism (SA) |
| Selection coefficients ($s$) | $s_{weak} = \{0.01, 0.02, 0.05, 0.1\}$<br>$s_{strong} = \{0.1, 0.2, 0.5, 1\}$ |
| Dominance coefficients ($h$, only defined for AP and SA) | 0.25<br>0.5 |
| Time of onset of selection ($T$, in generations) | **8Ne** |
| | 16$N_e$ |
| (Effective) population size ($N=N_e$) | 2,000<br>**20,000** |
| Recombination rate ($r$) | $10^{-7}$<br>**$10^{-8}$** |
| Mutation rate ($\mu$) | $10^{-7}$<br>**$10^{-8}$** |

To explore the effect of the strength of selection, equilibrium frequency and model of selection on the maintenance of the selected polymorphism (Figs. S1–S6) and the inference of balancing selection (Fig. 3), we ran simulations for a reference set with human-like parameters (bold values in Tab. 1, *N*_*e*_=20,000, µ = *r* = 10^-8^, *T* = 8*N*_*e*_ =160,000 generations) under OD, AP and SA across combinations of selection coefficients *s*_*i*_ and *s*_*j*_ drawn from two 4-by-4 grids with weaker selection {0.01,0.02,0.05,0.1} and stronger selection {0.1,0.2,0.5,1.0}. To explore the effects of mutation rate, recombination rate, selection onset time and *N*_*e*_, (Fig. 5), we ran similar sets of simulations for OD only, but changing the values of each of these parameters relative to the reference set of simulations (i.e. µ = 10^-7^, *r* = 10^-7^, *T* = 16*N*_*e*_ = 320,000 generations, *N*_*e*_ = 2,000).

In simulations to assess the maintenance of polymorphisms, we ran 100 replicate simulations in which the mutation successfully established (*i*.*e*., reached 5%) for each parameter combination. To assess the power to detect signatures of balancing selection, we complemented these with additional simulations to obtain sets of 100 replicate simulations for each parameter combination in which the polymorphism was maintained.

### Power of inference of balancing selection

We used the NCD (non-central deviation) method (Bitarello et al., 2018) to infer balancing selection on the simulated data (but for complementary results using BalLeRMix (Cheng & DeGiorgio, 2020), see Supplementary Material). Like most other current inference methods, NCD detects regions under balancing selection due to their signal of excess intermediate-frequency polymorphism. NCD for a genomic window with *n* informative sites is defined as:

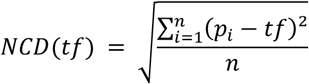

where *p*_*i*_ is the allele frequency at the *i*th informative site, and *tf* is the target frequency (set as an intermediate value *e*.*g*., 0.3, 0.4 or 0.5). The presence of a balanced site in the focal region with allele frequencies close to the target frequency then results in a low value of the NCD statistic, compared to a region under neutrality or purifying selection.

In our application of this approach, we used a refined version of the statistic, NCD2. This statistic gains power by also considering fixed differences (relative to an outgroup) in the site frequency spectrum (Bitarello et al., 2018), thus exploiting the signal of increased polymorphism and lower rate of substitution in regions under balancing selection. Although we have not simulated an outgroup, we can differentiate ancestral and derived alleles in our simulated genotypes and then manually add an ‘outgroup’ individual to the VCF file from our simulations, for which all sites are homozygous ancestral. Therefore, any sites that are fixed for the derived allele among the simulated (ingroup) individuals will be counted as fixed differences in the NCD2 calculation. We note that this approach adds less additional power than a real outgroup would, as the latter would also contribute fixed differences that arise due to substitutions in the outgroup lineage.

To assess the statistical significance of the NCD2 values, we generated a null distribution by running 1000 neutral simulations for each combination of relevant parameters (*N*_*e*_, µ and *r*) and calculating NCD2 for each. We then calculated a P-value for an NCD2 value from a simulation with selection by determining the proportion of NCD2 values from the null distribution (with corresponding *N*_*e*_, µ and *r*) that was lower than the NCD2 value with selection. Power was then calculated as the proportion of the 100 simulations generated under a particular set of parameters for which P ≤ 0.01.

### Statistical analysis of power estimates

We fitted binomial Generalised Linear Models (GLMs) to our power data to directly model the probability of detecting or failing to detect balancing selection as a function of the parameters that were varied in our simulations. The models used a logit link function and were fitted to the number of successful and unsuccessful instances of detection as the response variable. The significance of model terms was determined with an analysis of deviance, a series of sequential likelihood ratio tests based on the Chi^2^ distribution. This analysis further makes it possible to assess the importance of each model term through its deviance (the amount of variability in the response explained by a model term) relative to the null deviance, the total amount of variability in the response values analysed. We note that all models showed a degree of overdispersion, as indicated by the residual deviance exceeding the number of residual degrees of freedom several-fold. However, running alternative models that accommodate overdispersion (as a ‘quasibinomial’ model, combined with F-tests) produced qualitatively identical results.

These GLMs fit a linear (additive) model of detection probability (π) as a function of the simulation parameters. The model is fitted on the log-odds scale (where odds are *π*/(1 − *π*)) that accommodates the fact that probabilities are bounded by 0 and 1. The estimated model parameters, back-transformed to the odds-scale, provide information on the proportional change in the detection probability that is independent of the value of π (while the change in the actual probability due to a change in a predictor variable changes over the range of π).

## Acknowledgements

We would like to thank Filip Ruzicka and members of the Alliance EvolGen lab group for helpful discussions. MR, AA and DB were supported by grants from the UK Biotechnology and Biological Sciences Research Council, BBSRC (award BB/W007703/1 to MR and AA) and the Leverhulme Trust (award RPG-2021-414 to MR and AA). The work was initiated and developed during mutual visits between London and Melbourne by MR, AA and TC, made possible by an Australia Partnering award from the BBSRC (award BB/T019921/1 to MR and TC).

## Appendices

### Appendix 1 – Conditions for internal frequency equilibria

We model balancing selection making three simplifying assumptions that should apply broadly to many (perhaps most) cases where this type of selection is present. First, we focus on populations of diploid, randomly mating (outbred) individuals (see Glémin (2021) for a comprehensive treatment of balancing selection in self-fertilizing species). Second, we consider loci with two major allele types, which we label *A*_1_ (at a population frequency of *p*) and *A*_2_ (frequency 1 – *p*). Third, in scenarios involving antagonistic selection (i.e., antagonistic pleiotropy and sexual antagonism), we only consider scenarios of dominance that are amenable to balanced polymorphism, i.e. cases where each fitness component or sex either lacks dominance (fitness effects are additive in each context of selection) or there is a beneficial reversal of dominance, in which the costs of expressing the “wrong” allele for a given context of selection are partially or completely recessive (reviewed in Connallon and Chenoweth 2019; Grieshop et al. 2024). Our exposition draws upon previous studies of balancing selection theory (*e*.*g*., Robertson 1962; Kidwell et al. 1977; Curtsinger et al. 1994; Connallon and Clark 2013), with Table A1 providing an outline of the fitness values associated with each genotype and for each model of selection.

**Table A1.** Fitness per genotype for each model of selection.

|  | *Genotype* |  |  |
| --- | --- | --- | --- |
| | $A_1A_1$ | $A_1A_2$ | $A_2A_2$ |
| **Overdominance** | $1 - s_1$ | 1 | $1 - s_2$ |
| **Antagonistic pleiotropy** |  |  |  |
| **Fitness component 1** | 1 | $1 - t_1h$ | $1 - t_1$ |
| **Fitness component 2** | $1 - t_2$ | $1 - t_2h$ | 1 |
| **Overall Fitness** | $1 - t_2$ | $(1 - t_1h)(1 - t_2h)$ | $1 - t_1$ |
| **Sexual antagonism** |  |  |  |
| **Female fitness** | 1 | $1 - s_f h_f$ | $1 - s_f$ |
| **Male fitness** | $1 - s_m$ | $1 - s_m h_m$ | 1 |

Parameters for the three models fall within the following ranges: overdominance: 1 ≥ *s*_1_, *s*_2_ > 0; antagonistic pleiotropy: 1 ≥ *t*_1_, *t*_2_ > 0; sexual antagonism: 1 ≥ *s*_*f*_, *s*_*m*_ > 0. Dominance coefficients are within the range 0.5 ≥ *h, h*_*f*_, *h*_*m*_ ≥ 0, which includes additive effects (*h* = 0.5) and beneficial reversals of dominance (0.5 > *h*). Note that many of our analyses, we focus on scenarios where fitness effects are small, *i*.*e*..: 1 ≫ *s*_1_, *s*_2_ > 0; 1 ≫ *t*_1_, *t*_2_ > 0; and 1 ≫ *s*_*f*_, *s*_*m*_ > 0. Note that in the Appendices, we use slightly different notation to distinguish selection parameters of the OD, AP, and SA models (*s*_1_/*s*_2_, *t*_1_/*s*_2_, and *s*_*f*_/*s*_*m*_, respectively). For simplicity, and with no loss of generality, we use a single notation, *s*_1_/*s*_2_, in the main text.

#### Overdominant selection (OD)

With OD, the heterozygotes for the locus (*A*_1_*A*_2_) have the highest fitness and *A*_1_*A*_1_ and *A*_2_*A*_2_ homozygotes suffer fitness reductions of *s*_1_ and *s*_2_, respectively (Tab. A1). Evolutionary dynamics are captured by the following difference equation describing allele frequency change across a single generation:

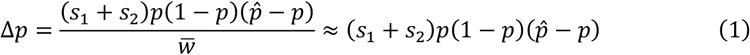

where 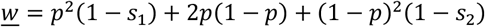 denotes mean fitness and:

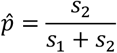

is the equilibrium frequency for the *A*_1_ allele. The final approximation in eq. (1) applies under weak selection (0 < *s*_1_, *s*_2_ << 1). Note that the entire range of parameter values in this model generates balancing selection (*i*.*e*., 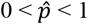 for the plausible parameter range, 0 < *s*_1_, *s*_2_ < 1).

#### Antagonistic pleiotropy (AP)

Following previous models (e.g. Curtsinger et al., 1994; Rose, 1982), we consider a locus with antagonistic pleiotropic effects on two major fitness components. Selection is directional within each component (Tab. A1) and fitness components combine multiplicatively to determine overall fitness. The evolutionary dynamics are described by:

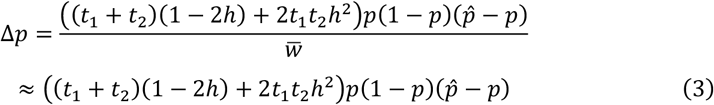

where 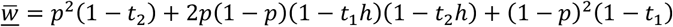 and the polymorphic equilibrium (whenever it is valid) is:

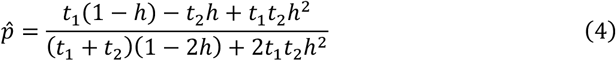

The equilibrium is valid under the following condition:

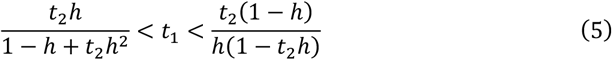

and otherwise, selection on total fitness is directional and favours fixation of either *A*_1_ or *A*_2_.

#### Sexually antagonistic selection (SA)

Following Kidwell et al. (1977), sexually antagonistic alleles are under opposing directional selection in males and females, characterised by a selection and dominance coefficient for each sex (Tab. A1). Letting *p* = (*p*_*f*_ + *p*_*m*_)/2 represent the overall frequency of *A*_1_ in the population, where *p*_*f*_ and *p*_*m*_ represent the respective frequencies in female and male gametes contributing to fertilization, the exact allele frequency dynamics are given by 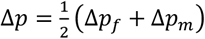, where:

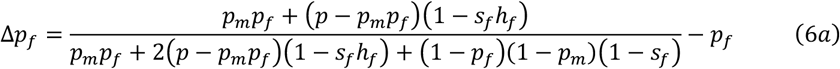

and

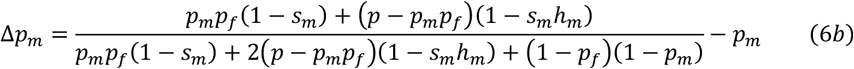

Within the range 0 ≤ *h* = *h*_*f*_ = *h*_*m*_ ≤ 0.5 which we consider here, and assuming modest to weak selection (0 < *s*_*f*_, *s*_*m*_ << 1), the allele frequency dynamics are closely approximated by:

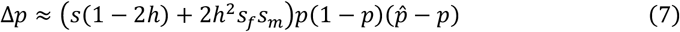

(Connallon and Clark 2013), where 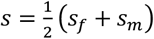, and 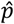 represents the polymorphic equilibrium when valid. With no dominance in either sex (*h* = 0.5), the equilibrium frequency (when it exists) is 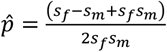, and for cases involving dominance reversals (*h* < 0.5), the equilibrium is 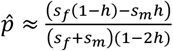. The condition favouring a polymorphic equilibrium (based on the exact eqs. (6a-6b)) is:

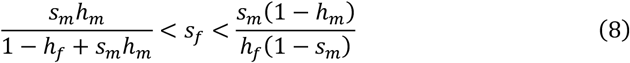

(Kidwell et al. 1977; Connallon and Chenoweth 2009).

If we assume that each selection coefficient is uniformly distributed with a mean of 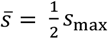 (where *s*_max_ represents the maximum selection coefficient, 0 < *s*_max_ ≤ 1), then the proportion of parameter space that results in balancing selection under AP will be:

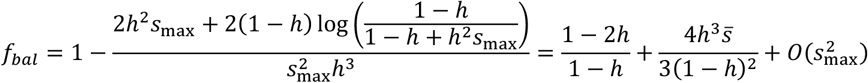

while that that results in balancing selection under SA is:

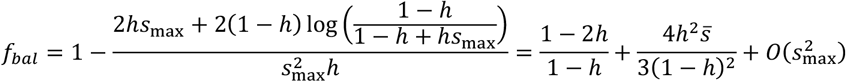

### Appendix 2 - Criteria for maintaining a long-term balanced polymorphism

A long-term polymorphism requires balancing selection to be both consistent over time and strong enough to offset the tendency of genetic drift to eliminate the polymorphism. Provided balancing selection is temporally consistent, then its capacity to maintain polymorphism can be quantified using standard diffusion theory (Rice, 2004; Wright, 1945). In the models presented above, the expected change in allele frequencies at a single locus under balancing selection (with change measured across a single generation) is:

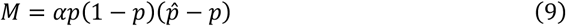

where *p* again refers to the frequency of *A*_1_, *p*^ is the equilibrium under balancing selection, and *α* represents a model-specific constant that reflects the strength of balancing selection (these can be extracted from eqs. (1), (3) and (7), which yield: *α* = *s*_1_ + *s*_2_ for the overdominance model, *α* = (*t*_1_ + *t*_2_)(1 − 2*h*) + 2*t*_1_*t*_2_*h*^2^ under antagonistic pleiotropy, and *α* = *s*(1 − 2*h*) + 2*s*_*f*_*s*_*m*_*h*^2^ under sexual antagonism). The variance in allele frequency change owing to drift is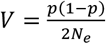, where *N*_*e*_ is the effective size of the population. Ignoring effects of recurrent mutation, the stationary distribution loci subject to balancing selection (*i*.*e*., the long-term probability density function for allele frequency *p*) is:

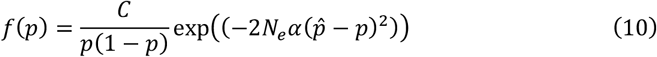

(Wright 1945; Robertson 1962; Rice 2004) where the constant *C* ensures that the probability density function integrates to one (*i*.*e*., 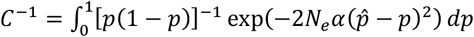.

Balancing selection effectively maintains polymorphism when the probability density near the equilibrium (*p* near 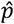) is high relative to the probability density near the boundaries of the distribution (*p* = 1/2*N*_*e*_ and *p* = 1 – 1/2*N*_*e*_). A measure of these relative densities (following Connallon & Clark, 2014) is given by:

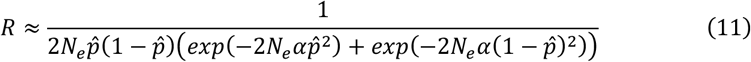

for which balancing selection is effective when *R* is large (*R* >> 1), and drift dominates when *R* is small (*R* < 1). Using a cut-off of *R* > 10 to represent effectively strong balancing selection, eq. (11) implies the following minimum values of *α* (*a*_min_) are required to ensure the stability of a balanced polymorphism:

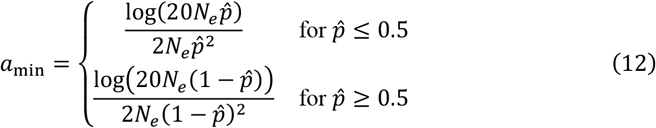

From eq. (12), balancing selection should be effective at maintaining polymorphism when *α* > *a*_min_, whereas drift will impede maintenance when *α* falls substantially below this threshold.

Figure 2 illustrates conditions for effective balancing selection across the spectrum of polymorphic equilibrium frequency states 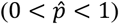 and for three different effective population sizes (*N*_*e*_ = 1,000; *N*_*e*_ = 10,000; *N*_*e*_ = 100,000). Two special cases of moderately strong balancing selection in a small population (*N*_*e*_ = 1,000 and *α* = 0.05) illustrate how the probability density of the stationary distribution is concentrated near 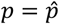 when *α* > *a*_min_, whereas it accumulates near *p* = 0 when *α* < *a*_min_ (Fig. 2B, top versus bottom panels).

### Appendix 3 - Linked variation

Long-term balanced polymorphisms can affect patterns of genetic diversity at linked and otherwise neutrally evolving loci, leading to characteristic signatures of inflated heterozygosity in genomic regions where the balanced polymorphism resides. With the exceptions of trans-species polymorphisms, it is these signatures that are used by current methods for the inference of balancing selection in genomic data (Bitarello et al., 2018; Cheng & DeGiorgio, 2020; Siewert & Voight, 2017). For a balanced polymorphism with alleles *A*_1_ and *A*_2_ that are stably maintained at equilibrium frequencies 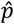 and 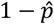 (respectively), the mean heterozygosity at a nearby neutral site is approximately:

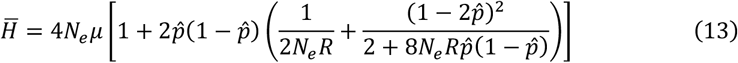

where *R* is the total recombinational distance between the neutral site and the site of the balanced polymorphism, and *μ* is the mutation rate at the neutral site (the approximation, which assumes that *R* >> *μ*, is obtained using mean coalescent times from p. 103 of Rice 2004).

The sum in the square brackets of eq. (13) determines the degree of inflation of heterozygosity at the linked site relative to that of an unlinked locus, where the expected heterozygosity would be 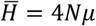. Recombination rate and the equilibrium frequency of the selected site also influence the width of the spike in neutral diversity that could be seen around a long-term balanced polymorphism. Let *R* = *nr* represent the cumulative recombination rate between the selected site and a given neutral site, where *n* is the number of nucleotide sites separating the two and *r* is the recombination rate per site (*i*.*e*., the probability of a recombination event between two directly adjacent nucleotide sites). Assuming that the polymorphism is intermediate 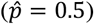, then the window around the selected site over which neutral diversity is elevated by at least two-fold (*i*.*e*.: where 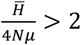) will be 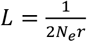 nucleotides, which implies that species with large effective population sizes and large per-nucleotide recombination rates should exhibit particularly narrow windows of inflated heterozygosity around loci maintaining long-term balanced polymorphisms.

## Notes

### Competing Interest Statement

The authors have declared no competing interest.

### Summary of Updates

Include supplementary material and reduce abstract to 250 words.

