## Supplementary Figures for "Characterising the detectable and invisible fractions of genomic loci under balancing selection"

Figure S1 - Maintenance of polymorphism under overdominance
Figure S2 - The effect of genetic drift on maintenance of polymorphism in simulations of overdominance
Figure S3 - Maintenance of polymorphism under antagonistic pleiotropy
Figure S4 - The effect of genetic drift on maintenance of polymorphism in simulations of antagonistic pleiotropy
Figure S5 - Maintenance of polymorphism under sexually antagonistic selection
Figure S6 - The effect of genetic drift on maintenance of polymorphism in simulations of sexually antagonistic selection
Figure S7 - Power to detect balancing selection in simulations of overdominance using NCD
Figure S8 - Power to detect balancing selection in simulations of antagonistic pleiotropy using NCD
Figure S9 - Power to detect balancing selection in simulations of sexual antagonism using NCD
Figure S10 - Power to detect balancing selection in simulations of overdominance using BalLeRMix
Figure S11 - Power to detect balancing selection in simulations of antagonistic pleiotropy using BalLeRMix
Figure S12 - Power to detect balancing selection in simulations of sexual antagonism using BalLeRMix
Figure S13 - Comparison of power of NCD and BalLeRMix across all parameter combinations presented in the main text.
Figure S14 - Power to detect balancing selection with NCD target frequency fixed at 0.5 across all simulated equilibrium frequencies
Figure S15 - Power to detect balancing selection with NCD as a function of NCD window sizes

**Figure S1 - Maintenance of polymorphism under overdominance**

Maintenance of polymorphism is 100% in our reference simulations (see Tab. 3 for parameter values). Maintenance is recorded as a proportion out of 100 initiated simulations with balancing selection. Panel A shows the grid of weak selection coefficients; panel B shows the grid of strong selection coefficients.

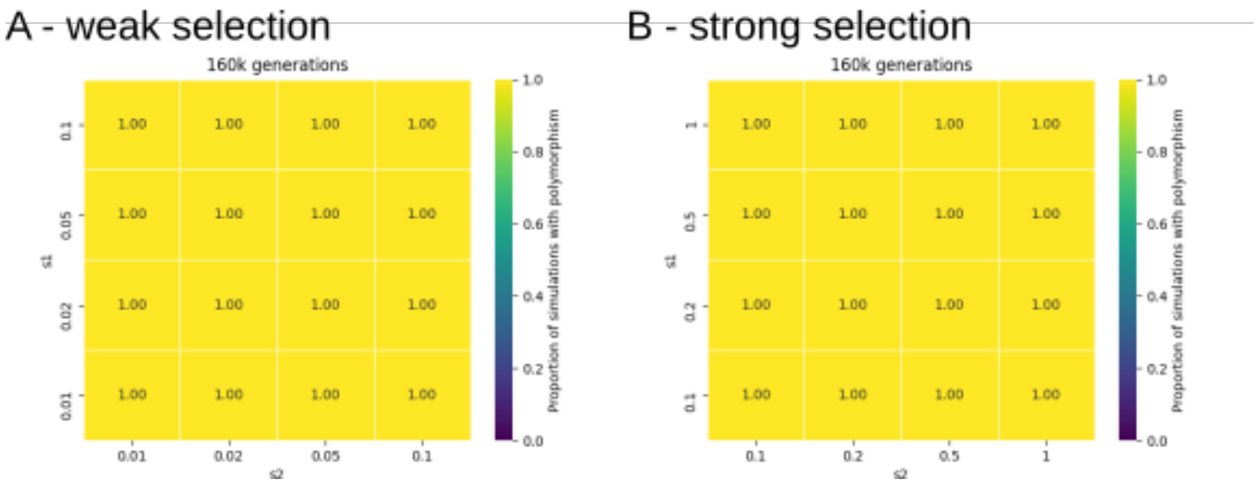

**Figure S2 - The effect of genetic drift on maintenance of polymorphism in simulations of overdominance**

Maintenance of polymorphism is reduced with increased genetic drift (i.e. lower effective population size ($N_e$), panels C and D, compared to higher $N_e$ in panels A and B), particularly at the edges of the parameter space, when selection coefficients are asymmetrical. Panels B and D show maintenance at $t=16N_e$ generations (twice as long as in the standard simulations, $t=8N_e$ generations). Maintenance is recorded as a proportion out of 100 initiated simulations with balancing selection. Simulation parameters not shown in the figure are: recombination and mutation rates $10^{-8}$.

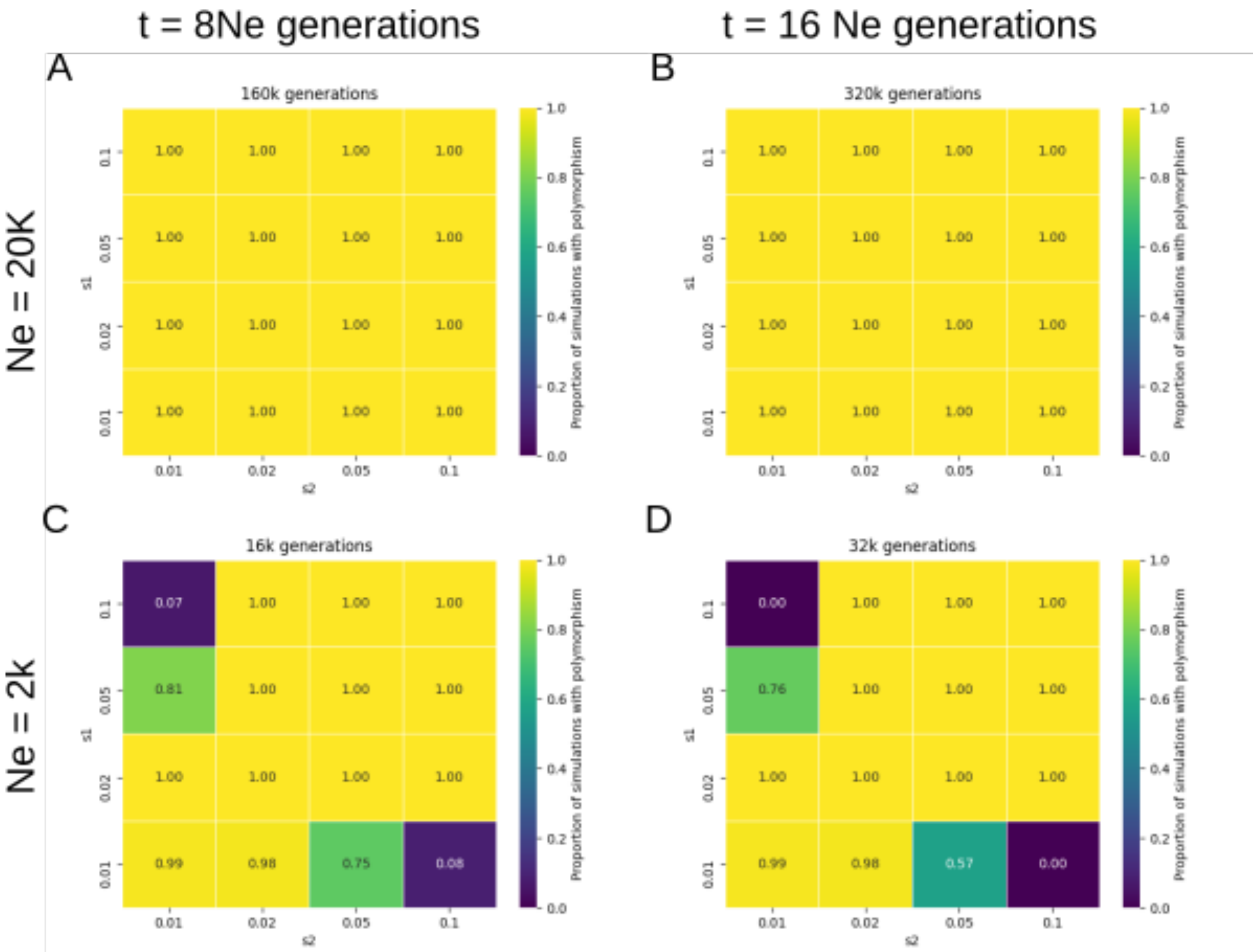

**Figure S3 - Maintenance of polymorphism under antagonistic pleiotropy**

Maintenance of polymorphism is lower at the edges of the parameter space, when selection coefficients are asymmetrical. Maintenance increases with dominance reversal (h=0.25, compared to codominance, h=0.5), and with stronger selection coefficients. Maintenance is recorded as a proportion out of 100 initiated reference simulations with balancing selection (see Tab. 3 for parameter values of reference simulations).

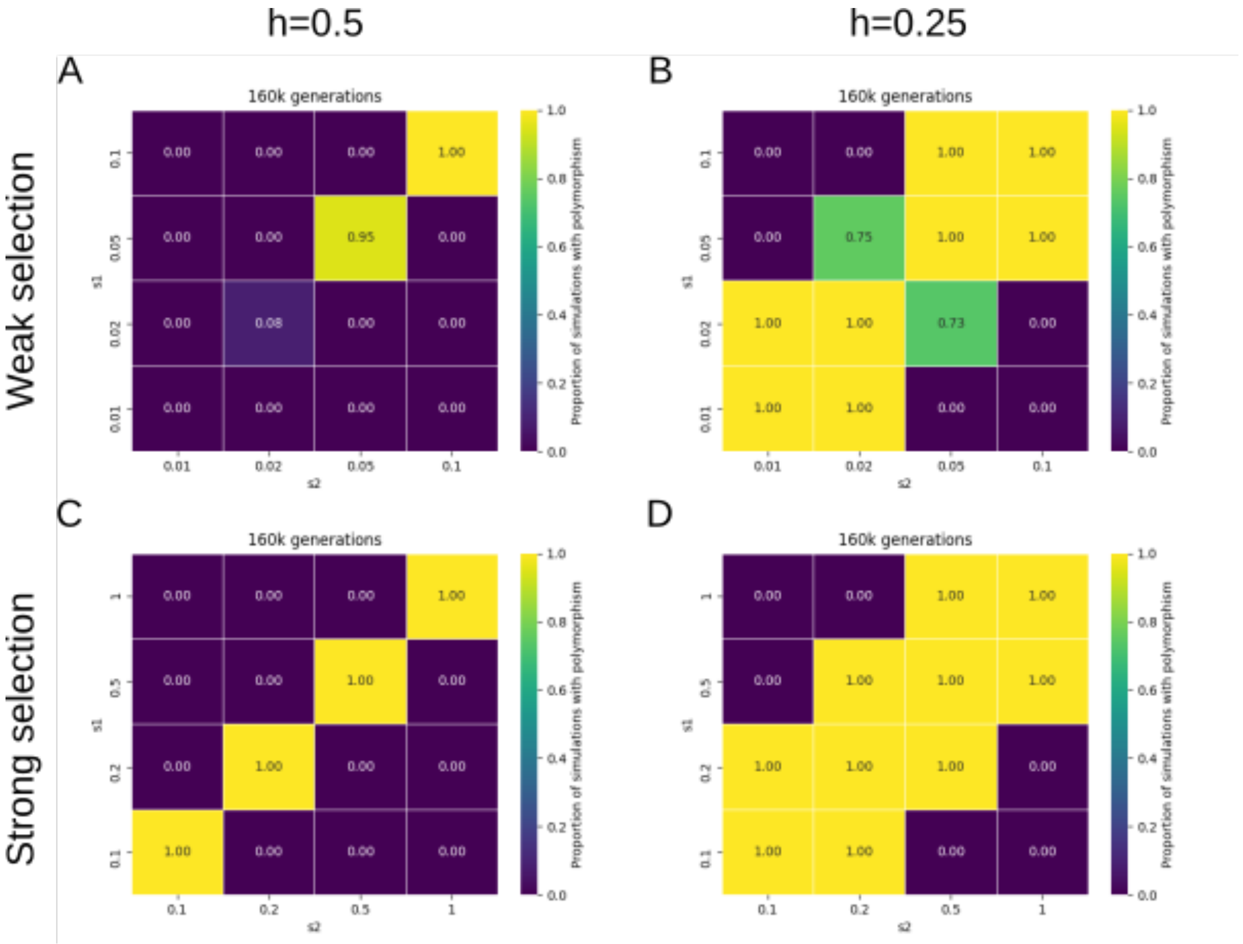

**Figure S4 - The effect of genetic drift on maintenance of polymorphism in simulations of antagonistic pleiotropy**

Maintenance of polymorphism is reduced with increased genetic drift (i.e. lower effective population size ($N_e$), panels A and B, compared to higher $N_e$ in Fig. S3 A and B), particularly at the edges of the parameter space, when selection coefficients are asymmetrical. Maintenance is recorded as a proportion out of 100 initiated simulations with balancing selection. Simulation parameters not shown in the figure are: recombination and mutation rates $10^{-8}$, selection onset time $t=8N_e$ generations.

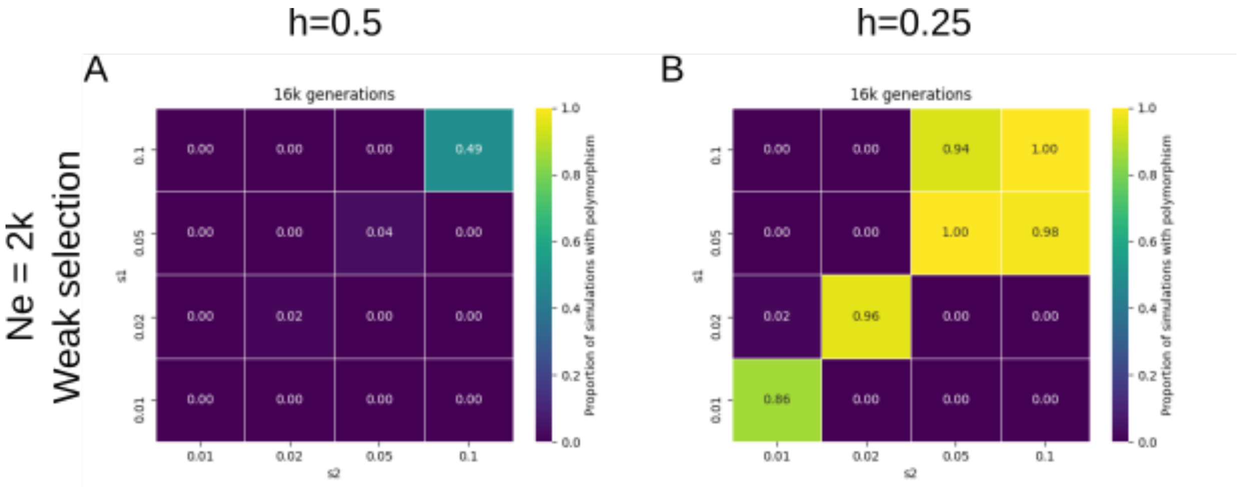

**Figure S5 - Maintenance of polymorphism under sexually antagonistic selection**

Maintenance of polymorphism is lower at the edges of the parameter space, when selection coefficients are asymmetrical. Maintenance increases with dominance reversal (h=0.25, compared to codominance, h=0.5), and with stronger selection coefficients. Maintenance is recorded as a proportion out of 100 initiated reference simulations with balancing selection (see Tab. 3 for parameter values of reference simulations).

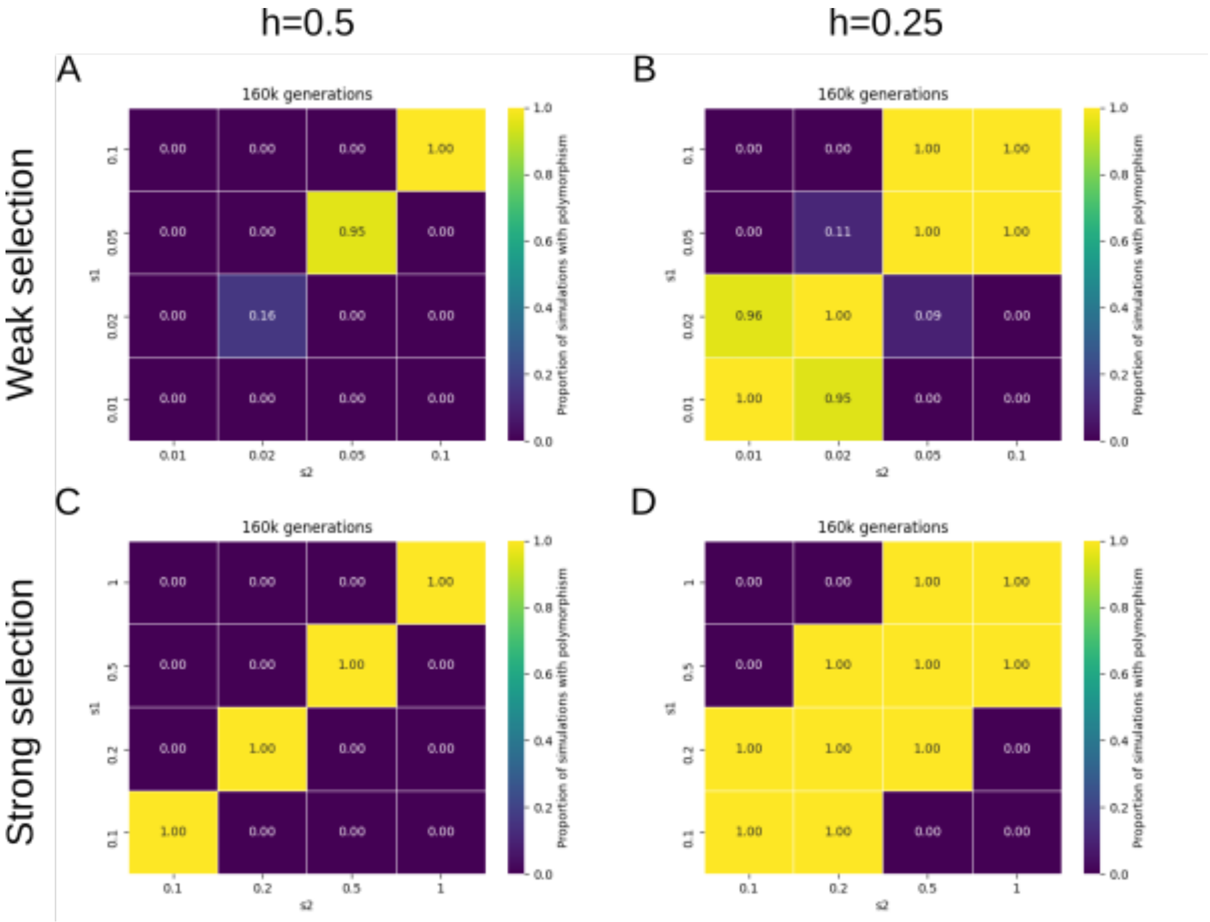

**Figure S6 - The effect of genetic drift on maintenance of polymorphism in simulations of sexually antagonistic selection**

Maintenance of polymorphism is reduced with increased genetic drift (i.e. lower effective population size ($N_e$), panels A and B, compared to higher $N_e$ in Fig. S5 A and B), particularly at the edges of the parameter space, when selection coefficients are asymmetrical and without dominance reversal (A, h=0.5). Maintenance is recorded as a proportion out of 100 initiated simulations with balancing selection. Simulation parameters not shown in the figure are: recombination and mutation rates $10^{-8}$, selection onset time $t=8N_e$ generations.

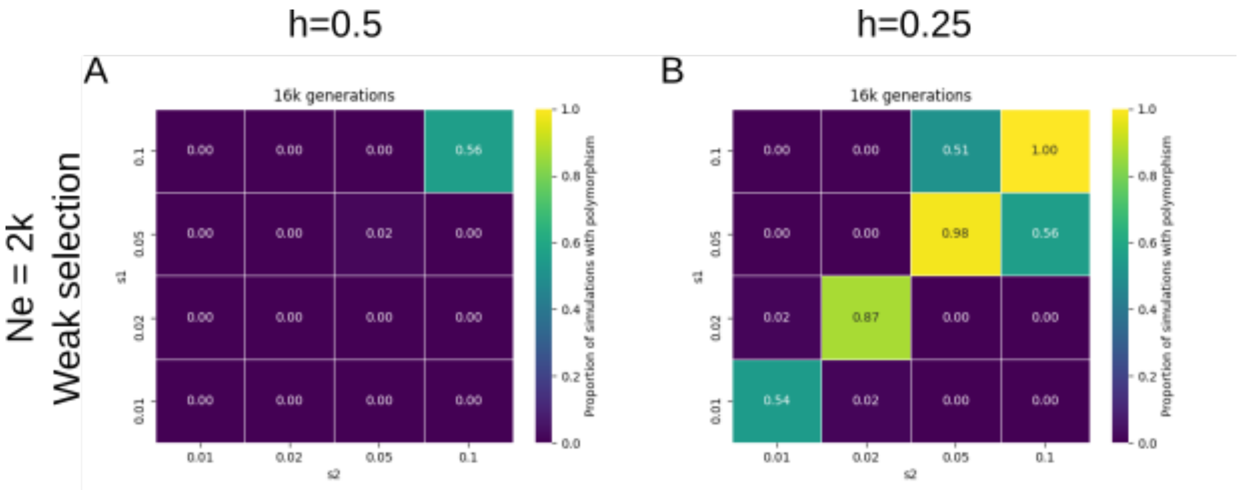

**Figure S7 - Power to detect balancing selection in simulations of overdominance using NCD**

Power is shown for reference simulations (parameter values are listed on the title of the plot).

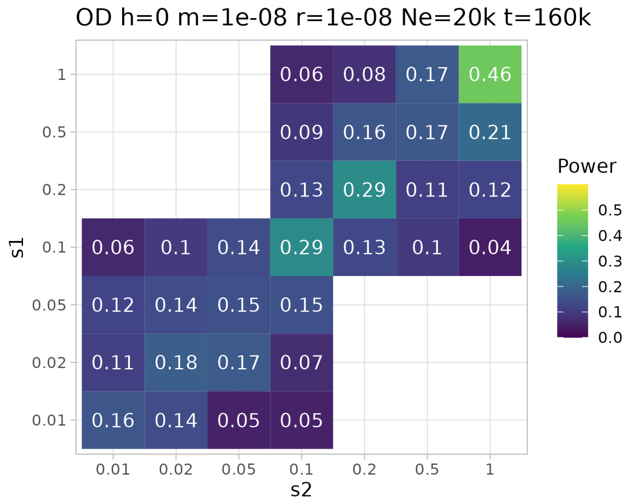


**Figure S8 - Power to detect balancing selection in simulations of antagonistic pleiotropy using NCD**

Power is shown for reference simulations (parameter values are listed on the title of the plots). A) Simulations under codominance (h=0.5); B) Simulations under dominance reversal (h=0.25).

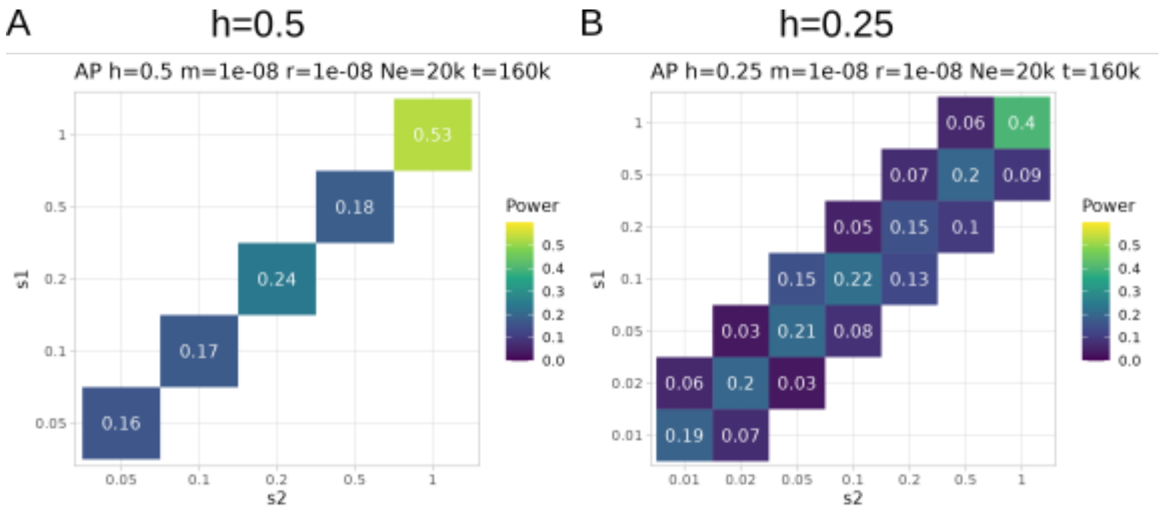

**Figure S9 - Power to detect balancing selection in simulations of sexual antagonism using NCD**

Power is shown for reference simulations (parameter values are listed on the title of the plots). A) Simulations under codominance (h=0.5); B) Simulations under dominance reversal (h=0.25).

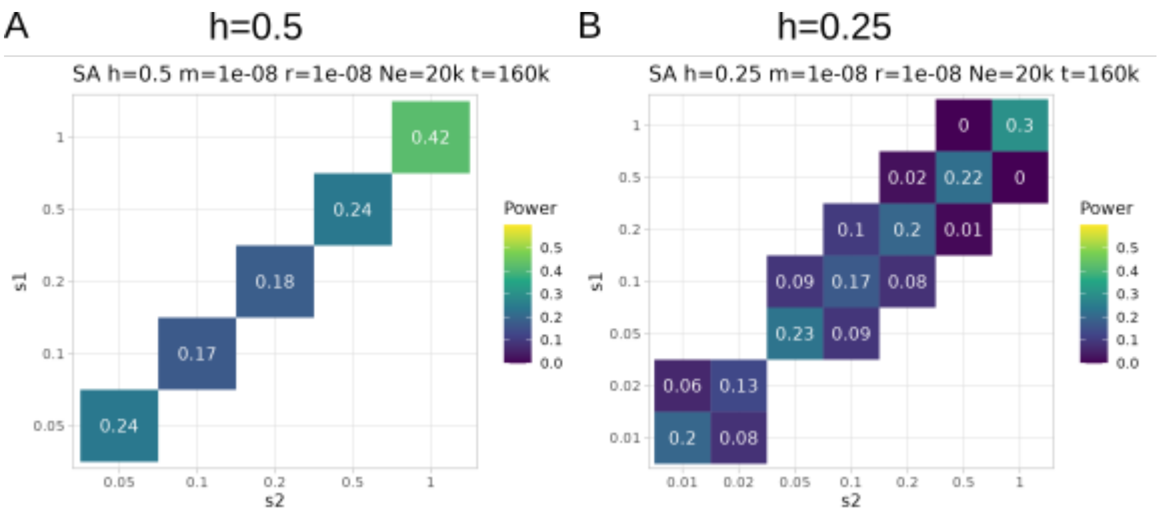


**Figure S10 - Power to detect balancing selection in simulations of overdominance using BalLeRMix**

Power is shown for reference simulations (parameter values are listed on the title of the plot).

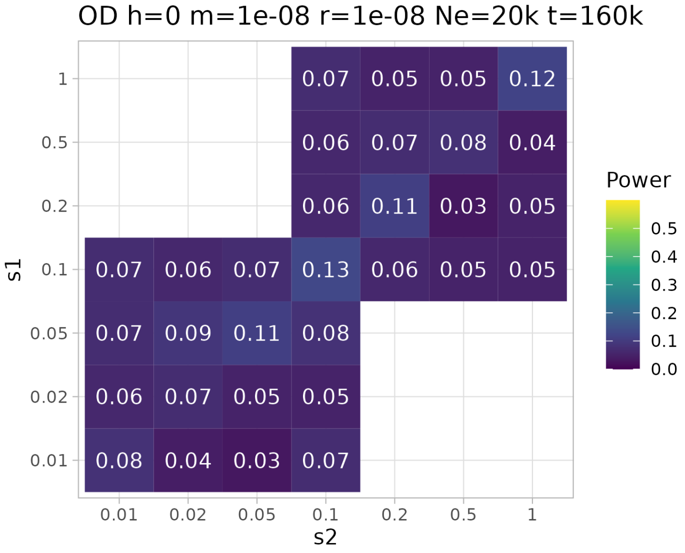

**Figure S11 - Power to detect balancing selection in simulations of antagonistic pleiotropy using BalLeRMix**

Power is shown for reference simulations (parameter values are listed on the title of the plots).

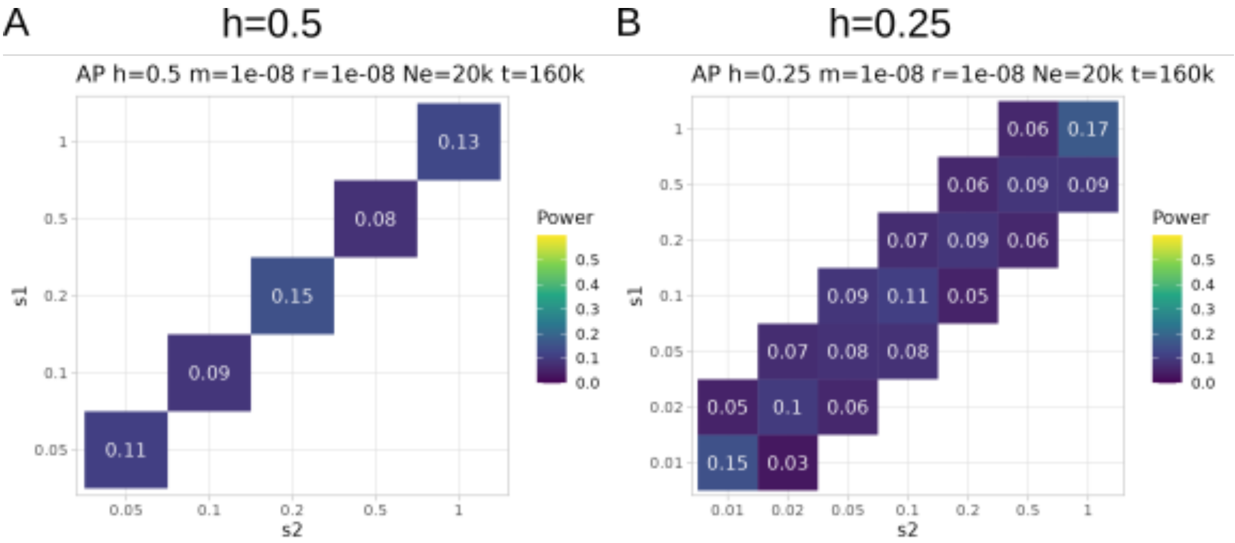


**Figure S12 - Power to detect balancing selection in simulations of sexual antagonism using BalLeRMix**

Power is shown for reference simulations (parameter values are listed on the title of the plots).

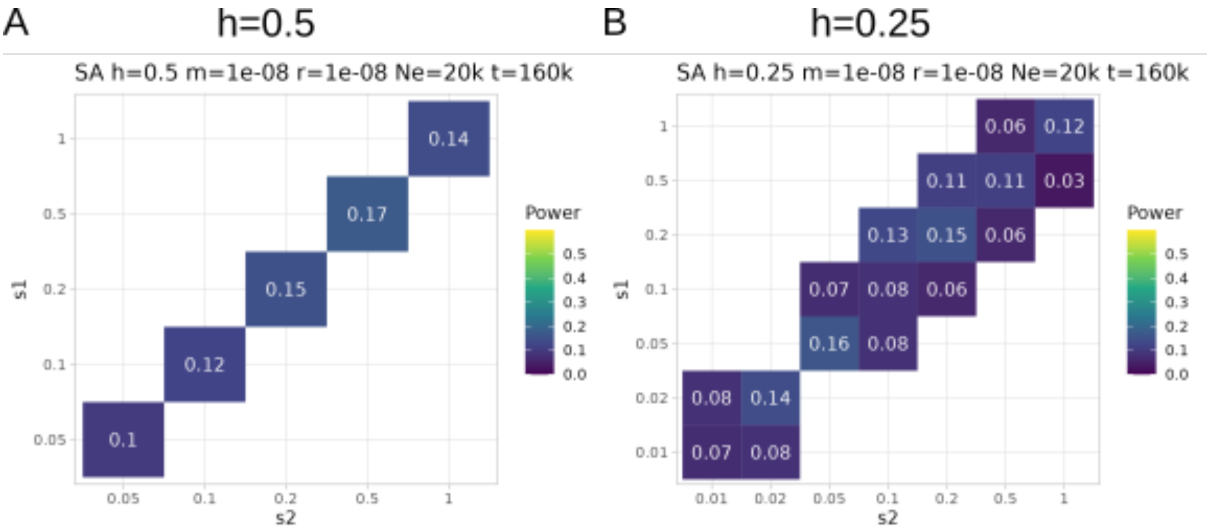

**Figure S13 - Comparison of power of NCD and BalLeRMix across all parameter combinations presented in the main text.**

Pearson correlation coefficient R=0.47, p-value<0.001. The diagonal line shows a slope of 1, corresponding to equal power of the two approaches.

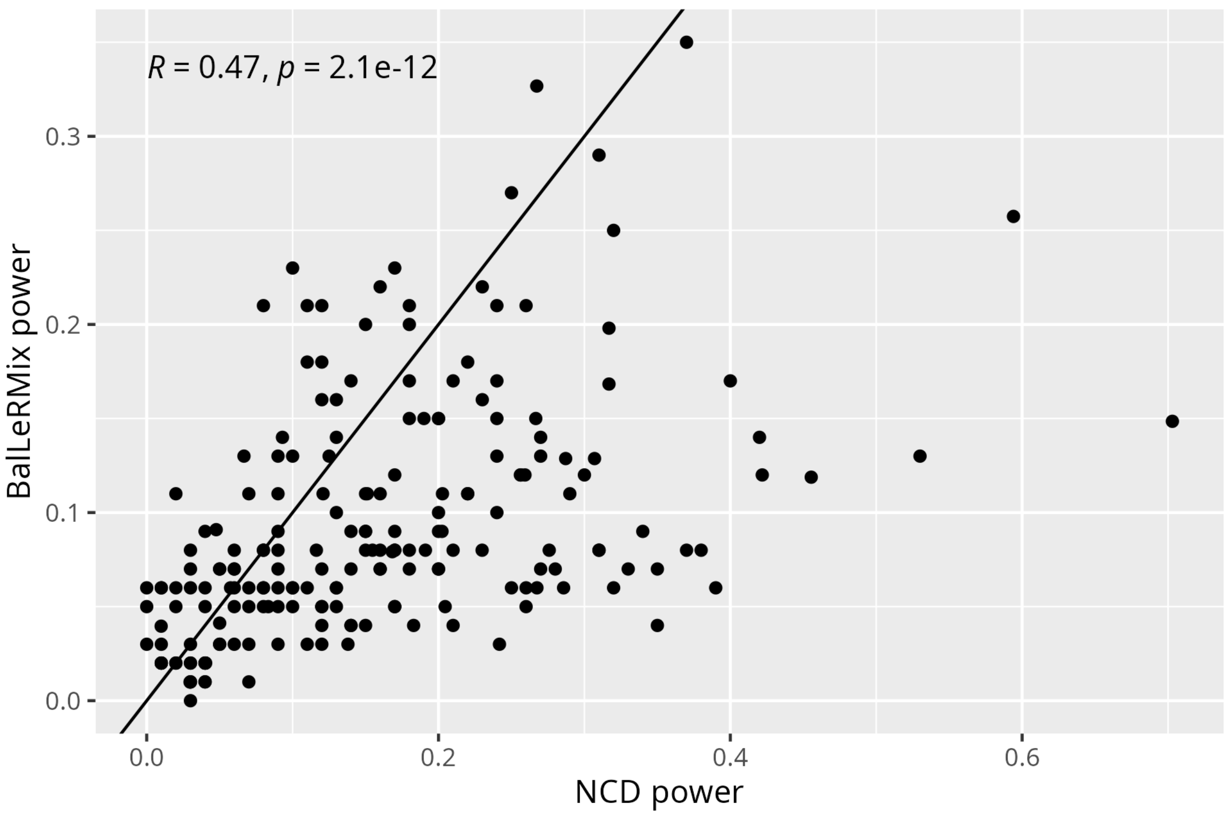

**Figure S14 - Power to detect balancing selection with NCD target frequency fixed at 0.5 across all simulated equilibrium frequencies**

Power is shown for data simulated under reference simulations of overdominance. A) NCD power with target frequency set as the true equilibrium frequency calculated from simulation parameters (same as Fig. S7). B) NCD power with target frequency fixed at p=0.5.

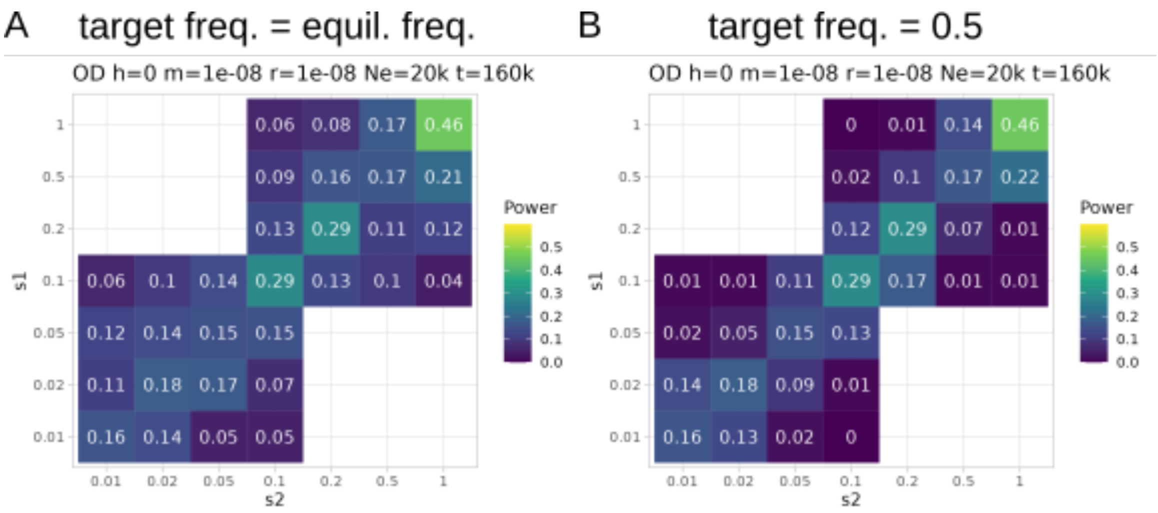


**Figure S15 - Power to detect balancing selection with NCD as a function of NCD window sizes**

Power was computed from data simulated under overdominance. Datapoints that are missing at smaller window sizes are due to small windows not having enough informative sites to compute NCD. The color legend shows the equilibrium frequency (here indicated as p*). A) Reference simulations. B) Simulations with increased recombination rate.

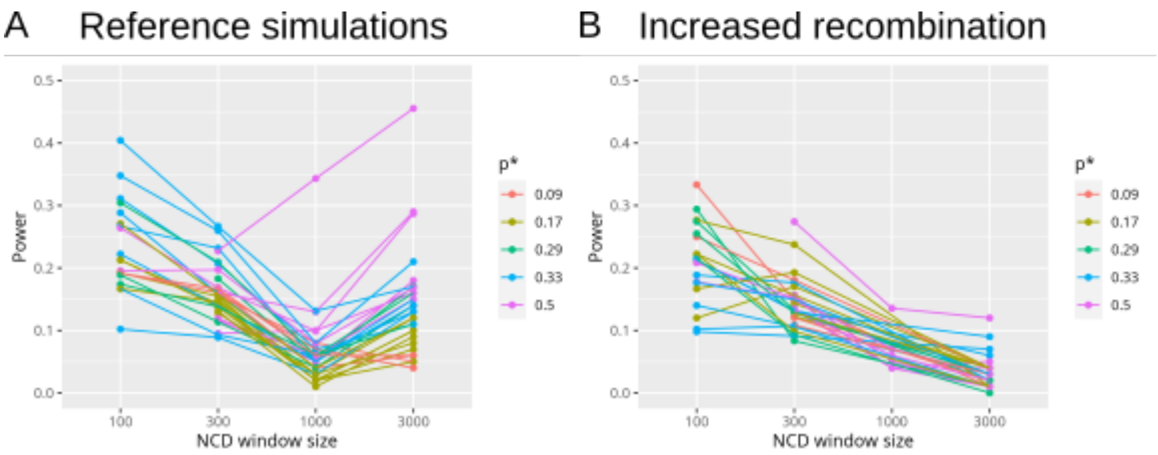
